# Revisiting the Out of Africa event with a novel Deep Learning approach

**DOI:** 10.1101/2020.12.10.419069

**Authors:** Francesco Montinaro, Vasili Pankratov, Burak Yelmen, Luca Pagani, Mayukh Mondal

**Author notes:** Contributed Equally (Francesco Montinaro, Vasili Pankratov). Correspondence and requests for materials should be addressed MM.

## Abstract

Anatomically modern humans evolved around 300 thousand years ago in Africa^1^. Modern humans started to appear in the fossil record outside of Africa about 100 thousand years ago though other hominins existed throughout Eurasia much earlier^2–4^. Recently, several researchers argued in favour of a single out of Africa event for modern humans based on whole-genome sequences analyses^5–7^. However, the single out of Africa model is in contrast with some of the findings from fossil records, which supports two out of Africa^8,9^, and uniparental data, which proposes back to Africa movement^10,11^. Here, we used a novel deep learning approach coupled with Approximate Bayesian Computation and Sequential Monte Carlo to revisit these hypotheses from the whole genome sequence perspective. Our results support the back to Africa model over other alternatives. We estimated that there are two successive splits between Africa and out of African populations happening around 60-80 thousand years ago and separated by 12-13 thousand years. One of the populations resulting from the more recent split has to a large extent replaced the older West African population while the other one has founded the out of Africa populations.

## Introduction

In the last few decades, the development of efficient and powerful computing infrastructure allowed us to gain substantial progress in the machine learning field, especially for computationally demanding algorithms such as Neural Network (NN)^12,13^ and Bayesian methods^14,15^. NN was demonstrated to be an useful tool for specific types of tasks, such as classification or natural language processing^12–14,16^. However, NN requires a large amount of data as a training set. In some cases, simulated datasets are one of the strategies to overcome this limitation. The simulation of synthetic genetic data can be helpful to substantially mitigate this problem^17,18^. NN is already adopted in population genomics studies to interpret the genomics data in terms of underlying demography^19–21^ and positive selection^22,23^. However, unlike classical approaches, it is still challenging to measure the significance of a prediction performed by NN, given that it is a black-box approach. Approximate Bayesian Computation (ABC) can be used to weigh the accuracy of a NN-based prediction from the data itself, without knowing the maximum likelihood function^19,20,24^.

Recent fossil record analysis suggests that anatomically modern humans appeared around 300 thousand years ago (kya) in Africa^1^. This hypothesis is corroborated by genetic data^25^, which projected the deepest splits between modern human populations at a similar time interval. Although fossil records advocate that there might be multiple Out Of Africa (OOA) events for modern humans^26^, recent genetic studies revealed that all modern non-African or OOA populations fit a model characterised by a single OOA event^5–7^. This conclusion indicates that older OOA migrations, documented by archaeological records, might have not left much contribution to modern human populations, with the possible exception in Oceania (Papuan populations)^27^ and some archaic hominin^28^.

While the single OOA model finds support in both autosomal and uniparental data^11,29,30^, there is some evidence for a more complicated scenario. Most of the uniparental haplogroups are closer to each other in OOA populations than African haplogroups (thus having less time to the most recent common ancestor [TMRCA]), corroborating a single clean OOA model, apart from the sister Y haplogroups D and E. The haplogroup D can be found in isolated populations in Asia (i.e., Andamanese, Tibetan, Japanese, etc.), while the haplogroup E is ubiquitous in sub-Saharan African populations. They are slightly closer to each other than any other haplogroups found in OOA populations from them^11,31^. This observation might be explained by a back to Africa migration^11^ or a more complicated scenario^32^. Some autosomal analyses also suggest that the separation between Africa and OOA populations might not be a single split event^33–36^.

Testing these hypotheses (single out of Africa, back to Africa and two out of Africa) is challenging due to the strong bottleneck of non-African populations^37–39^, differential archaic introgression between populations^5,19,40^ and various migrations within Africa^36,41^. The lack of ancient genomic data older than 15 kya^42^ from Africa or the Middle East makes it difficult to address this issue from an ancient DNA perspective. However, NN have been shown to be extremely powerful to disentangle such complex scenarios^19^. Here, we present ABC-DLS (Approximate Bayesian Computation using Deep Learning and Sequential Monte Carlo method) which allows us to infer the most likely scenario among different competing demographic models as well as to estimate their parameter values with high precision. Our approach relies on a NN trained on simulated genetic data under the models being tested. However, it has three key improvements compared to other similar approaches. First, the use of the hdf5^43^ data format and tensor flow^44,45^ allows for extremely large training datasets. Second, the conventional NN approach is augmented using ABC which helps to provide statistical support for the NN prediction and to obtain posterior distribution for the model parameter values. Third, inspired by previous works^46^, we applied a modification of the Sequential Monte Carlo (SMC, also known as the Particle Filter method)^47^ approach to iterate the whole procedure. This improved the accuracy substantially compared to previously implemented methods^19,48^. We apply this method to test the three OOA models mentioned above.

## Results

### ABC-DLS

The general workflow for ABC-DLS (both for model selection and parameters estimation) includes the following steps. First, we simulated^18^ multiple sets of genetic data for each tested model using demographic parameters sampled from a uniform distribution within prior ranges (Table 1). Next, we converted this data into joint site frequency spectrum (SFS) (although potentially any other summary statistics (SS) can be used) and split the data into a training and a testing subset. We then trained the NN (implemented using TensorFlow^44^ with Keras backended^45^) on the training dataset to either select between demographic models or to estimate the demographic parameters. The resulting NN is applied to the testing dataset as well as to the observed SS data (see below as well as Methods for more details). Next, we apply ABC to estimate support for the NN prediction on the observed data comparing the NN prediction outcome between the observed data and the testing dataset (see Methods, Supplementary Figure 2 and also our previous paper ^19^). Finally, in cases when SMC is used, we essentially iterate the parameter estimation step by SMC. When estimating the posterior range for the parameters using ABC, we kept the top five percent (equal to the tolerance level) of simulations from the testing dataset that best matched with the observed data. We then used the parameters of those simulations to update our prior range and sent it for next iteration till convergence reached (Supplementary Figures 2 and 3).

**Table 1:** Prior parameters range used for producing the Site Frequency Spectrum (SFS).

| Parameters | OOA_S | OOA_B | OOA_M |
| --- | --- | --- | --- |
| N_A | 5,000 - 25,000 | 5,000 - 25,000 | 5,000 - 25,000 |
| N_AF | 10,000 - 150,000 | 10,000 - 150,000 | 10,000 - 150,000 |
| N_EU | 10,000 - 150,000 | 10,000 - 150,000 | 10,000 - 15,0000 |
| N_AS | 10,000 - 150,000 | 10,000 - 150,000 | 10,000 - 150,000 |
| N_F | 5,000 - 30,000 | 5,000 - 30,000 | 5,000 - 30,000 |
| N_EU0 | 500 - 5,000 | 500 - 5,000 | 500 - 5,000 |
| N_AS0 | 500 - 5,000 | 500 - 5,000 | 500 - 5,000 |
| N_B | 500 - 5,000 | 500 - 5,000 | 500 - 5,000 |
| N_BC | NA | 500 - 30,000 | NA |
| N_AF0 | NA | 500 - 30,000 | NA |
| N_MX | NA | NA | 500 - 30,000 |
| N_B0 | NA | NA | 500 - 30,000 |
| T_FM (ky) | 2 - 5 | 2 - 5 | 2 - 5 |
| T_FS (ky) | 0.1 - 10 | 0.1 - 10 | 0.1 - 10 |
| T_DM (ky) | 10 - 50 | 10 - 50 | 10 - 50 |
| T_EU_AS (ky) | 10 - 30 | 10 - 30 | 10 - 30 |
| T_NM (ky) | 5 - 50 | 5 - 50 | 5 - 50 |
| T_XM (ky) | 5 - 50 | 5 - 50 | 5 - 50 |
| T_Mix (ky) | NA | 5 - 50 | 5 - 50 |
| T_Sep (ky) | NA | 5 - 50 | 5 - 50 |
| T_B (ky) | 5 - 270 | 5 - 220 | 5 - 220 |
| T_AF (ky) | 5 - 700 | 5 - 700 | 5 - 700 |
| T_N_D (ky) | 330 - 450 | 330 - 450 | 330 - 450 |
| T_H_A (ky) | 120 - 250 | 120 - 250 | 120 - 250 |
| T_H_X (ky) | 450 - 700 | 450 - 700 | 450 - 700 |
| NMix (%) | 1 - 3 | 1 - 3 | 1 - 3 |
| DMix (%) | 0 - 2 | 0 - 2 | 0 - 2 |
| XMix (%) | 0 - 10 | 0 - 10 | 0 - 10 |
| FMix (%) | 0 - 10 | 0 - 10 | 0 - 10 |

NA means not applicable. Ky means kilo or thousand years.

Before testing our primary hypothesis on real sequence data, we tested if our new approach (ABC-DLS) is robust enough for the known results. The predicted parameters for real sequence data (see later for more details) are consistent with previous works from the literature^37,39,49^(Supplementary Table 1). We also simulated models (model S, B, M, see later for more information) and created mock observed SS (simulation parameters coming from Table 2, Supplementary Table 2 and 3). We found that our novel approach with SMC predicted the right model for every case, suggesting it can find the correct model.

**Table 2:** Posterior range for parameters of model B.

| Parameters | Mean | CI | Events (kya) |
| --- | --- | --- | --- |
| N_A | 13,500 | 13,495 - 13,501 |  |
| N_AF | 21,689 | 21,277 - 22,496 |  |
| N_EU | 119,988 | 111,986 - 134,779 |  |
| N_AS | 122,321 | 114,667 - 135,974 |  |
| N_F | 20,137 | 17,149 - 24,731 |  |
| N_EU0 | 1,813 | 1,766 - 1,871 |  |
| N_AS0 | 730 | 719 - 752 |  |
| N_BC | 20,347 | 13,024 - 25,099 |  |
| N_B | 2,123 | 2,100 - 2,164 |  |
| N_AF0 | 28,491 | 27,967 - 28,758 |  |
| T_FM (ky) | 3.4 | 2.1 - 4.9 | 3.4 (2.1 - 4.9) |
| T_FS (ky) | 4.9 | 0.2 - 9.7 | 8.3 (3.4 - 13.2) |
| T_DM (ky) | 15 | 14.4 - 15.8 | 15 (14.4 - 15.8) |
| T_EU_AS (ky) | 17.7 | 17.2 - 18.3 | 32.7 (31.8 - 33.6) |
| T_NM (ky) | 7.2 | 6.7 - 7.4 | 39.9 (38.9 - 40.9) |
| T_XM (ky) | 14.7 | 13.7 - 15.7 | 47.5 (46.1 - 48.8) |
| T_Mix (ky) | 15 | 14.1 - 16.1 | 47.7 (46.4 - 49.1) |
| T_Sep (ky) | 11.2 | 10.8 - 12.3 | 59 (57.4 - 60.5) |
| T_B (ky) | 13.2 | 12.7 - 13.5 | 72.2 (70.6 - 73.7) |
| T_AF (ky) | 208.9 | 196.8 - 218.3 | 281.1 (270.2 - 291.9) |
| T_N_D (ky) | 447.2 | 444.3 - 448.9 | 447.2 (444.3 - 448.9) |
| T_H_A (ky) | 249.4 | 247.2 - 250.5 | 696.6 (693.8 - 699.5) |
| T_H_X (ky) | 695.4 | 686.1 - 700.3 | 695.4 (686.1 - 700.3) |
| Mix (%) | 91.98 | 89.66 - 93.08 |  |
| NMix (%) | 3.01 | 2.98 - 3.02 |  |
| DMix (%) | 0.63 | 0.58 - 0.68 |  |
| XMix (%) | 5.04 | 4.85 - 5.14 |  |
| FMix (%) | 2.37 | 2.13 - 2.49 |  |

CI is the confidence interval of 2.5%-97.5% of respective parameters. Ky means kilo years and kya means kilo or thousand years ago from now.

### Model Selection

To test our hypothesis, we simulated three OOA models: Simple model (model S), Back to Africa model (model B), and Mix model (model M) with all the models having introgression from Neanderthal to all OOA populations^50^, Denisova or Unknown to Asia^19,51,52^, African Archaic to Africa^36,53,54^ and European Farmers to Africa^55^ (NDXF) (see methods for more details, Supplementary Figure 1 and Table 1). We used HGDP dataset^56^ of five Yoruba (African), five French (European) and five Han Chinese (East Asian) as our real dataset. Next, we used three different methods to choose between the competing models: i) ABC-RF that combines random forests with ABC (here onwards referred to as RF)^48^; ii) NN and ABC together (here onwards referred to as DL) which is analogous to our previously published method ABC-DL^19^; and iii) the novel method introduced here ABC-DLS which augments the DL method with SMC (here onwards referred as DLS). Although all three methods identified the model B as the most probable one, the prediction certainty varied between methods (Table 3). While DLS returned 100% probability for model B, DL and RF gave lower support. Also, when 10 independent runs were tested, model B won 10 times out of 10 using DLS and 9 out of 10 using DL. Moreover, Bayes factor value was predicted to be 6.69 between model B and model S by DL. These suggest that we cannot reject model S completely with DL. This difference in prediction certainty was likely due to the better power of DLS to differentiate between the three models compared to the other (Table 3).

**Table 3:** Cross validation and Model Selection using different approaches.

| **RF** | OOA_B | OOA_M | OOA_S |
| --- | --- | --- | --- |
| OOA_B | 95.33% | 1.23% | 3.44% |
| OOA_M | 1.06% | 85.12% | 13.82% |
| OOA_S | 4.45% | 8.94% | 86.61% |
| Posterior of votes | 70.40% | 13.20% | 16.40% |
| **DL** |  |  |  |
| OOA_B | 97.36% | 0.53% | 2.12% |
| OOA_M | 0.03% | 85.74% | 14.23% |
| OOA_S | 1.09% | 10.55% | 88.36% |
| Posterior model probabilities | 87.27% | 0.13% | 12.6% |
| **DLS** |  |  |  |
| OOA_B | 99.84% | 0.04% | 0.13% |
| OOA_M | 0.00% | 100.00% | 0.00% |
| OOA_S | 0.00% | 0.08% | 99.92% |
| Posterior model probabilities | 100.00% | 0.00% | 0.00% |

Confusion matrix for misclassification is reported here using RF (Random Forest), DL (only Neural Network) and DLS (Neural Network and Sequential Monte Carlo together) for random samples from the models with ABC. Posterior of votes and Posterior model probabilities are final posterior after using the real data.

The DLS results were reproduced under different data filtering strategies and different datasets (Supplementary Table 6). As our base models assumed four pulse migration events based on previous studies (three introgression scenarios and recent migration of Neolithic farmers), we tested if these assumptions could affect our inference. We tested different models with 1) No introgression and no farming migration (NI), 2) Neanderthal and Denisova introgression (ND), 3) Neanderthal, Denisova and Africa Archaic introgression (NDX) 4) Neanderthals, Denisova introgression with farming migration (NDF) using only DLS. Except for the no introgression model (Supplementary Table 7), we always found model B to be supported over models S and M. When we compared all these 15 models together ([B, M, S] x [NI, ND, NDX, NDF, NDXF]) using DLS, model B with Neanderthal, Denisova, African archaic introgression, and Neolithic migration (BNDXF) is supported over all other possibilities (P(BNDXF|data) =0.76) (Supplementary Table 8). This result not only demonstrated the robustness of our inference for model B but also independently supported other assumptions which were reported before but not all of them were confirmed together^19,36,50–52,54,55^. We would also like to point out a simpler model without Neolithic migration (P(BNDX|data=0.24) cannot be rejected by our approach.

### Parameter Estimation

After demonstrating that model B best explains the real SS data, we used the three methods described above (RF, DL and DLS) to estimate the model’s parameters. The confidence intervals returned by DLS are much narrower than those of the alternative approaches (Table 2, Supplementary Tables 9 and 10) and comparable with other methods^37,39,49^ thus showing good performance of our new method. Hence, all the results discussed below are the ones obtained with DLS.

Our inference suggests that there was first a separation between the Ancient African population (AA) and a population ancestral to both Back-to-Africa and the actual Out-of-Africa populations (OOA’) around 72.2 (CI 70.6 - 73.7) kya followed by a split between back to Africa (B2A) and OOA 59 (CI 57.4 - 60.5) kya and an admixture between AA and B2A 47.7 (CI 46.4 - 49.1) kya. The Neanderthal introgression to OOA happened much later, 39.9 (CI 38.9 - 40.9) kya, suggesting that this Back to Africa migration cannot explain the Neanderthal ancestry found in modern African populations^55^. Our method predicted the admixture proportion from B2A to be as high as 92% (CI 89.66 - 93.08) suggesting a massive replacement of the AA population.

Our results also comply with Y-chromosomal phylogeny and support back to Africa as proposed before^11^. However, our estimation time of separation between populations is much younger than what is reported in Y-chromosomes. One explanation might be that we used a slightly higher mutation rate (1.45×10^-8^ per bp per generation)^57^ instead of a slightly slower alternative (1.25×10^-8^ per bp per generation)^58,59^. When we used the slower mutation rate, our estimation for most of the events time increased (Supplementary Table 11). Indeed, the separation time between B2A and OOA populations corresponds to 67 (CI 66.3 - 67.6) kya, which is close to the estimate of TMRCA between Haplogroup D and E (72 kya^11^).

To independently validate our results, we compared effective population size (Ne) trajectories and cross-coalescent rates obtained by applying Relate^32^ to real data as well as to data simulated under each of the three models using the mean posterior parameters (Table 2 and Supplementary Table 2 and 3) predicted by DLS (following a flowchart represented in Supplementary Figure 3a)^34^. We observe a close match between the estimates for the real data and our best predicted model (Figure 2) which suggests our parameter estimation to be accurate. This similarity is particularly interesting, given that we have not used any LD-based SS to optimize those parameters. On the other hand, neither the Ne trajectory nor the cross-coalescent rate over time is informative to differentiate between the three models (data not shown). Specifically, the gradual separation between African and OOA populations, which was observed before with Relate and similar methods^33,34^, cannot be directly explained by the back to Africa or two out of Africa migration as this separation is also matched in our model S (Supplementary Figure 4).

**Figure 1:**
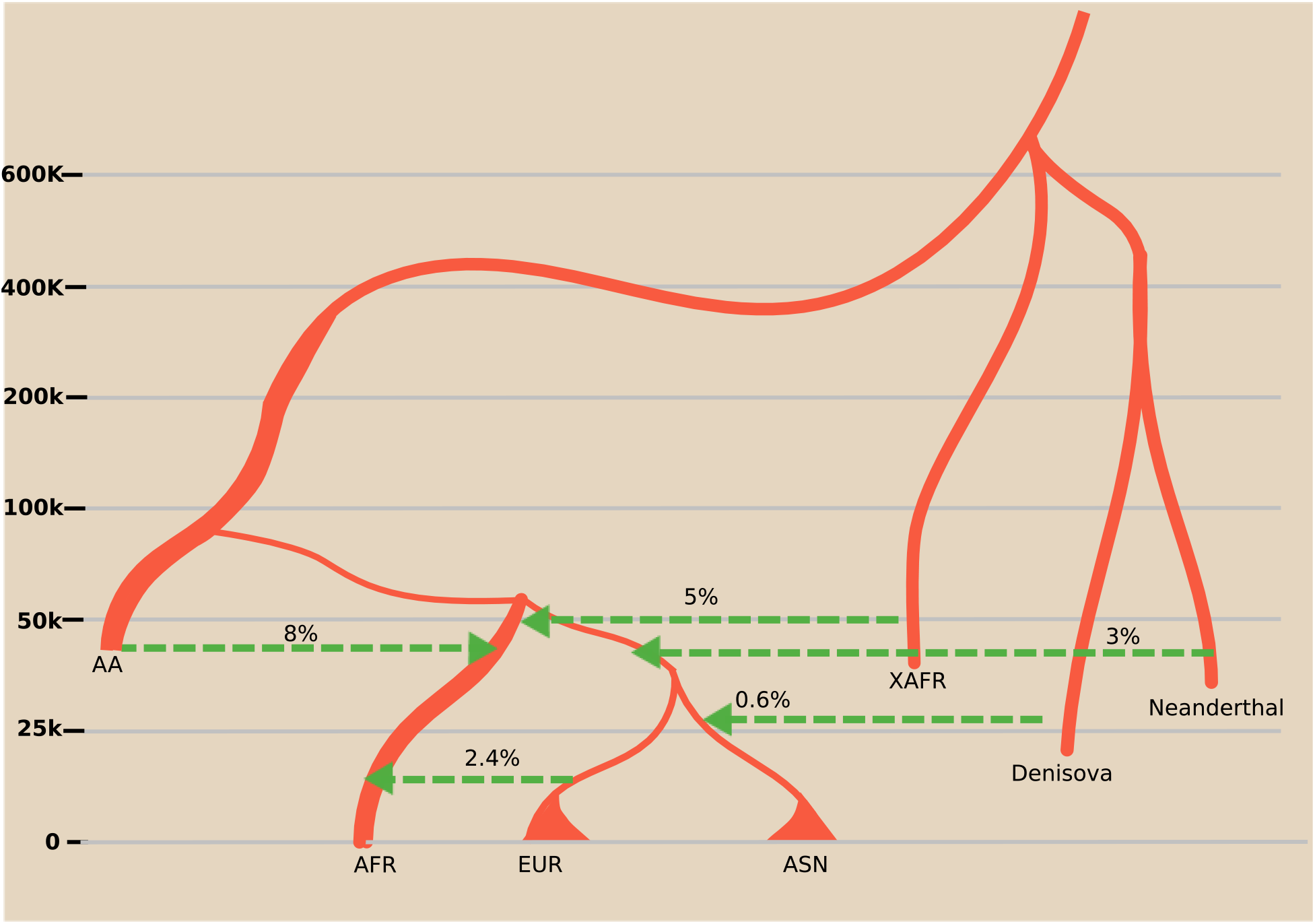
Schematic of Inferred Demography. Model B with only mean posterior. Kya is kilo years ago, AFR is African, EUR is Europeans, ASN is East Asian, NEAN is Neanderthal, DENI is Denisova and XAFR is African Archaic.

**Figure 2:**
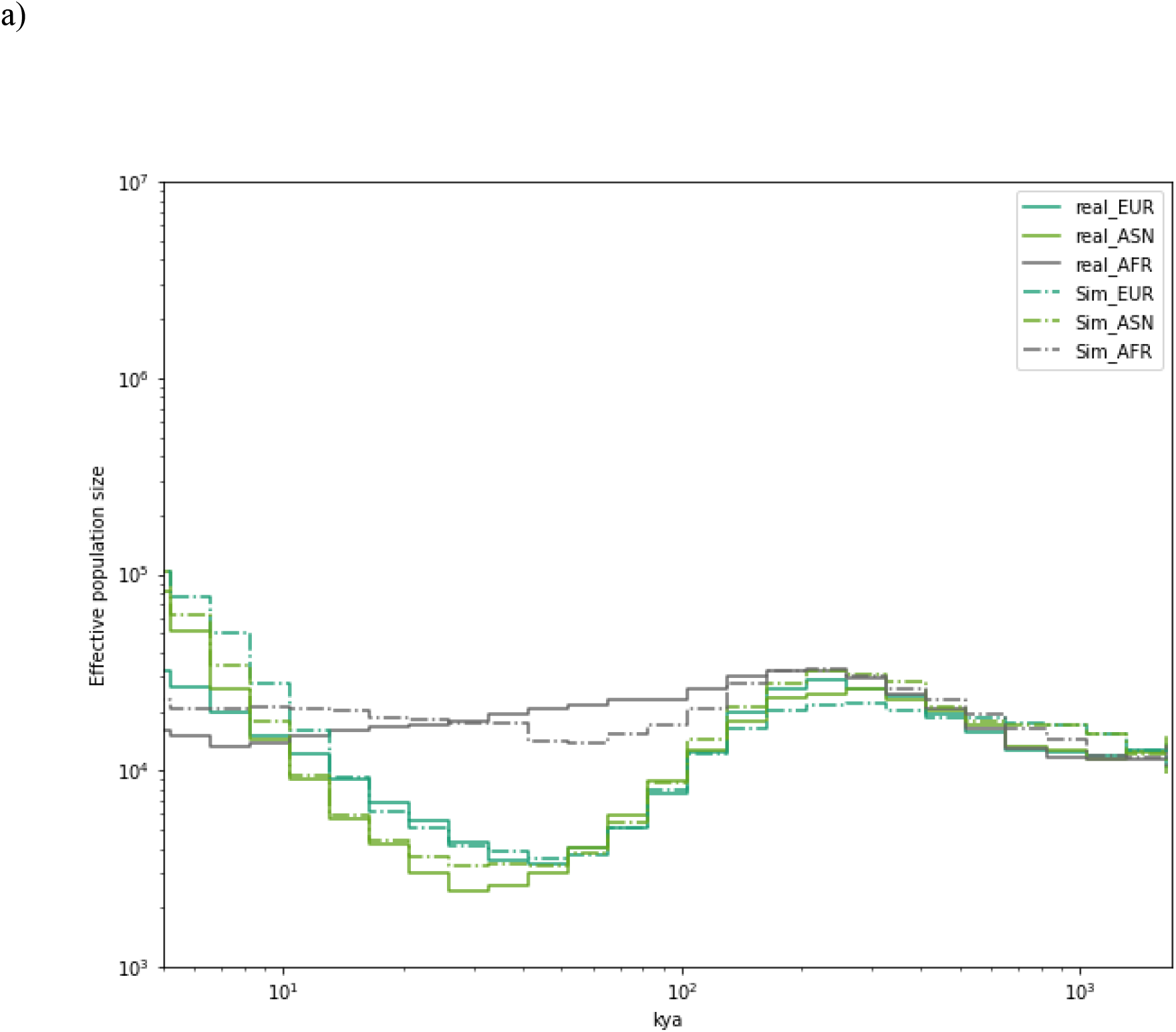
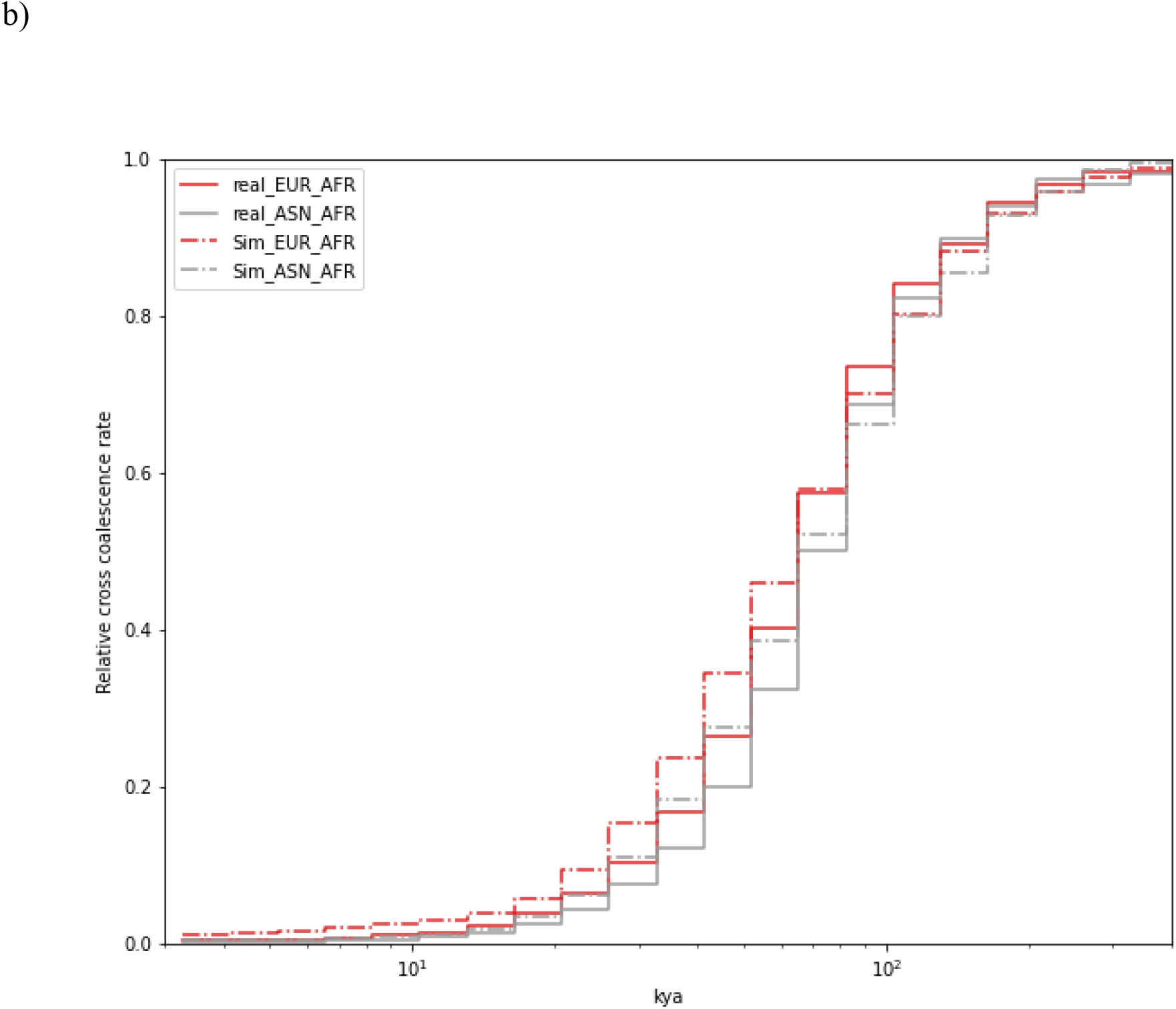
Effective population size trajectories and coalescence rates over time. Here we compare effective population size (a) and relative cross-coalescence rates (b) estimated using Relate between the real and data simulated under the model B. a) x axis in kya (kilo years ago) and y axis is the effective population size for corresponding populations presented in the inset. Both axes are in log scale. b) x axis is in kya and y-axis shows the relative cross coalescent rate for corresponding populations pairs presented in the insert. x axis is log scale.

## Discussion

We presented here that the ABC analysis can be substantially improved by using NN coupled with the SMC approach. Our methodology is robust to test any hypotheses which can be simulated, which cannot be extensively tested by other methods (especially for scenarios of admixture from ghost populations where the ancient genomes are unavailable) and can accommodate any kind of SS. In this study, we used SFS as SS because it is effortless to calculate and have sufficient information^37,60^. Our results might be further improved by using some LD-based SS^53,61^ but we opted out as they are computationally demanding to produce and the improvement in the result is minimal (at least for the tested scenario). Although our approach (DLS) is fast enough, the main bottleneck currently is the production of the simulated SS data.

In our models, we have not adopted any migration rates between populations, although our approach can use it. This is because we found out that our approach (Parameter Estimation using DLS) predicted non-zero migration rates when we used a mock observed SS data coming from a pulse model with no migration (mean values from Table 1) and a NN trained on an island model with migrations (Supplementary Table 4 and 5). This suggests that models including migration rates may lead to equifinality as suggested by others^62^ and/or our approach is incapable of estimating them.

Although in our scenarios model B is preferred over model S, considering no introgression as an option (NI) supported model M over other models (Supplementary Table 7). This result might be a side effect of the Neanderthal introgression in OOA. Under certain conditions (i.e., older separation time between Africa and OOA [T_B]), model M with no introgression and model S with Neanderthal introgression are comparable (Neanderthal population behaves like the first OOA population in this scenario). This result suggests a possible drawback of our method as different demographic histories can give similar SFS patterns, which can bias our interpretation if not incorporated in the model correctly^63^ and also advocates for the importance of parameter estimation as it can give insight for the choice of model selected.

Although the estimated proportions of introgression from Archaic populations have values consistent with those previously reported^28,53^, the separation time between *Homo sapiens* and archaic populations are more recent than those previously inferred^50,64^ if we used a loose prior of 400-1,100 kya. These deviations were not reproduced when we used simulated SS generated under known parameters from Table 2. This may be specific to real sequence data and might be a side effect of some of our assumptions (for example some unknown interactions between these populations which was not modelled here) or systematic biases due to the use of European reference genome^65^ or recent changes of generation time or mutation rate per generation^66,67^. Thus, the admixture with archaic populations may be seen as a way of introducing noise in the simulations for model selection rather than an attempt to obtain true parameter estimates. Most probably in the future, we can improve this estimate by directly using the available ancient genomes together with modern datasets.

We cannot also reject a simpler model of no Neolithic migration^55^. Even if we assume the Neolithic migration affected Yoruba, the predicted total length of Neanderthal sequence in an average Yoruba genome would be less than 5 Mb compared to the 17 Mb identified by Chen et al^55^. This discrepancy also cannot be explained by the back to Africa model as introgression happened much later after the separation. This suggests that most of the Neanderthal signal in Yoruba should be explained by some other migration (for example from Human to Neaderthal^28^).

Our results suggested a back to Africa model (model B) is more likely than a simple out of Africa event (model S). Although this model is better in explaining the real data, it might not be the final one. An even more complicated migration or admixture model which was not tested here might still better explain the real data. We have not tested two out of Africa events directly, although our model M is similar to two out of Africa model under certain conditions (assuming that European and East Asian do not have differential admixture with first OOA population). It will be interesting to revisit this hypothesis with Papuan populations in the future.

We would like to caution that although we are naming the model “Back to Africa”, the OOA population did not need to be geographically out of Africa^68^. Our estimates, particularly the effective population size of B2A (N_BC) and the time of Neanderthal introgression (T_NIntro), advocate that the split might have happened within Africa itself before the actual out of Africa event. In such a case, our results can be explained by the separation of West and East African population 80 kya (T_B) and then later the primary separation of OOA and East African population 67 kya (T_Sep) (assuming mutation rate of 1.25×10^-8^ per bp per generation^58,59^ and generation time of 29 years^69^). In this regard, our model is more akin to Lipson et al. 2020^36^ model rather than what is suggested by Cole et al. 2020^35^. If we assume model from Lipson et al. to be true, the most parsimonious explanation would be that our B2A population represents Basal West African population which separated from OOA populations 67 kya (T_Sep). Our AA represents Ghost modern^36^ which contributed to modern West African population around 10% which admixed around 60 kya from our prediction. On the other hand, if we assume true back to Africa, then most likely the OOA event took place less than 80 kya (T_B). This suggests that most of the older fossils (>80 kya) found outside Africa^2–4^ are unlikely to have contributed to OOA populations (assuming the ancestor of all modern human originated in Africa and never left Africa before OOA event). Geographical location where B2A separated from OOA is immensely important for this hypothesis but cannot be estimated from our approach. It will be especially fascinating to test this hypothesis using ancient genomes from those areas from that time point when they will be available.

## Methods

### Real Data

We have downloaded the high coverage HGDP vcf files^56^ and randomly selected five African (Yoruba, YRI), five European (French, FRN), and five East Asian (Han Chinese, HAN) individuals. As an alternative dataset, we have also downloaded the high coverage 1000 Genome^70^ vcf files with personal communication from Michael Zody of New York Genome Center (this data is yet to be published). We randomly selected five individuals from Africa (Yoruba from Ibadan, Nigeria YRI), Europe (Utah Residents with European origin, CEU), and East Asia (Chinese Han from Beijing, CHB) population. We kept both the data set separated and kept only positions present in every individual (within every data set), bi-allelic and Single Nucleotide Polymorphism (SNP). We lifted the genome to GR37 using Picard tools. Moreover, we filtered out regions with genes and CpG islands (for more details, please see Mondal et al. 2019^19^). Independently, we also used a mappability mask for the HGDP dataset in Supplementary Table 6. All the filterings were done with combinations of vcftools and bcftools^71,72^. The vcf file was converted to SFS using an in-house code using scikit allel^73^.

### Simulations

All the simulations were done in msprime^18^. We have produced the joint site frequency spectrum (SFS) of five individuals per populations (African, European, and East Asian genome) simulating one mega base pair (Mbp) of replicates with the recombination rate of 10^-8^ per base pair (bp) per generation and the mutation rate of 1.45×10^-8^ per bp per generation^57^. We also alternatively used 1.25×10^-8^ per bp per generation for mutation rate (only for Supplementary Table 11)^58,59^. Here, we kept the recombination rate constant, as SFS is not affected by the local recombination rate^63^. We assumed generation time of 29 years^69^.

In msprime, Admixtures were represented as MassMigration (the fraction of a population replaced by another population in a single generation). In contrast, migration rates under island models (where appliable) were represented as Migrationrate (the rate of fraction per generation of a population was replaced by another population for several generations).

The ABC-DLS analysis is efficient enough to be done on a single computer. The main bottleneck of the whole approach is the production of the SFS data. Msprime is fast, but the total amount of data, which needs to be simulated for the NN, is impossible to produce in a single computer. We have used a snakemake pipeline to produce the SFS on the cluster^74^.

### Demographic models

#### Simple out of Africa (model S)

In this simulation model, we have modeled a simple OOA event (Supplementary Figure 1) closely following Gravel et al.^39^, except the migration rates are assumed to be nil. When we simulated models with migrations rates, we slightly modified the model proposed by Gravel et al^39^. Migration rates are denoted by m_pop1_pop2, where pop1 is the population that received the migration, and pop2 is the population from where the migration originated. If the migration rates are bi-directional and equal, we have four parameters (Supplementary Table 1) like in Gravel et al^39^. However, if they are not equal, we have eight parameters (Supplementary Table 4 and 5) to have all the combinations between African, European, East Asian and OOA population.

#### Back to Africa (model B)

In this model, the basic OOA model still holds plus additional changes required for Back to Africa migration (Supplementary Figure 1) are added. The basic idea is drawn from Poznik et al.11. In this scenario, the OOA’ population is separated into populations B2A and OOA T_Sep generations ago before the separation between European and Asian populations (which happens between T_B and T_EU_AS generations ago). Next, B2A migrated to Africa having an effective population size of N_BC and mixed with the African population (AA) T_Mix generations ago with a mixing proportion of Mix (the portion of AA ancestry replaced). After the admixture, the effective population size of the African population is changed from N_AF0 to N_AF.

#### Mixed out of Africa (model M)

This model (Supplementary Figure 1) is similar to model S with an additional population M separating from the African population around T_Sep generation ago and having an effective population size of N_MX. M mixed with OOA at T_Mix generations ago with Mix being the proportion of OOA ancestry being replaced by M. After the admixture, the effective population size of OOA is changed from N_B0 to N_B. The basic idea came from Haber et al^32^ as well as two OOA^27^.

#### Other Migrations as Prior

We also added some pulse migrations or admixtures proposed by different studies on top of these basic models. We simulated OOA to have introgression from Neanderthal^50^ around T_NM generation ago with the proportion of NMix. After the separation between Europeans and East Asians, the East Asian population has introgression from Denisova^51,52^ or an unknown population^19^ around T_DM and the amount is DMix. Neanderthal separated from Denisova or the unknown around T_N_D generation ago, and Neanderthal-Denisovan lineage separated from the modern human lineage T_H_A generation ago^28,64^. The African population also has introgression from another unknown archaic population^36,53,54^, which introgressed around T_XM generations ago with the proportion of XMix. This unknown population separated from modern human lineage around T_H_X generation ago. We found out that our method is incapable of finding the effective population size for archaic populations. Thus, we assumed them equal to N_A same with the ancestral effective population size. We also simulated Neolithic farmers, which separated from Europeans around T_FS generations ago with effective population size of N_F and admixed with the African population around T_FM generation ago with the proportion of Fmix^55^.

Some events can only happen after a particular event has already taken place (for example, the separation of European and Asian populations can only happen after the Neanderthal introgression, based on our prior assumption). The relations between these events are not straightforward and written in Supplementary Table 12.

#### Genotype to Site Frequency Spectrum

We ran several simulations and converted every instance of simulations into a joint multidimensional unfolded site frequency spectrum (SFS) of a three-dimensional array from three populations: Africa, Europe, and East Asia. SFS is the total number of segregating sites for a given derived allele count present in each population.

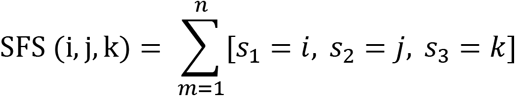

Where, i, j, k= The number of derived alleles count per SNP in pop1, pop2 and pop3 respectively. n= the total number of segregating SNPs.

The SFS was generated from simulations by msprime^18^ and then represented in a row. A similar conversion was done on the real data from the vcf file. Although it is possible to use any SS for our approach, we only used SFS as our choice of SS, given that it is straightforward to obtain and informative enough^37,60^. All the elements of real SFS were multiplied by a constant (frac) to make it comparable with the length of simulated regions if they do not match. For example, we multiplied the real SFS by 10 / 647 if we simulate a 10 Mbp region per simulation, and the real data is coming from 647 Mbp region (after filtering).

### ABC-DLS

We have used TensorFlow with Keras backend^44^ for building the NN and used a simpler version of the SMC approach^47^ to improve the prediction.

#### Parameter Estimation with DL

Here we describe parameter estimation using NN with ABC. We ran a total of 60,000 different simulations, with every simulation producing 3,000 of 1 Mbp regions (3 Gbp [giga base pair] in total, roughly equal to the length of the human genome). Throughout all our steps, we always simulated regions of 1 Mbp replicates, as they are fast to produce. Every line is one such simulation performed under a given demographic model with the first few columns being the parameters used for that simulation and the rest of the columns representing SFS elements. We ran Parameter estimation on this CSV file to retrieve the parameters predicted on observed data for the given model. We used the known parameters as labels for training the NN (y), and we used the SFS as input (x). Thus, we can think of NN here as an inverse function of the simulation. We kept 10,000 random lines for the testing dataset and ABC analysis, and the rest were used for training the NN. All the columns of SFS and parameters were normalized with MinMax scaler^75^, so the whole data is within 0.0 and 1.0 per column.

We used four hidden layers of a dense NN (Supplementary Figure 5) with activation relu, and we used linear for the output layer with the same number of units as the number of parameters. We used a Masking layer at the beginning to remove cells with zero from the learning algorithm and then Gaussian noise injection of .05 to introduce some noise (Supplementary Figure 5). We used logcosh as a loss function and Nadam for the optimizer. The NN ran through the training dataset several times (epochs) to increase accuracy. We used EarlyStopping on loss coming from validation dataset with the patient of 100 and used the ModelCheckpoint of the lowest loss result on validation data. We also used ReduceLROnPlateau of factor 0.2 to reduce the learning rate if we reached minima for several epochs (10 by default).

After training is done, we used the testing dataset to predict the parameter from the SFS, which was then used for cross-validation tests and parameter prediction using loclinear from ABC^76^ with the tolerance of 0.01.

This approach is similar to what we have published before in ABC-DL^19^, although we have used the latest tools (TensorFlow, Keras, and hdf5). The whole approach is presented in a flowchart (Supplementary Figure 2).

#### Model Selection with DL

Here we describe model selection using NN and ABC. After running simulations for the three demographic models (sample numbers are same as above per model), we produced three corresponding CSV files. These CSV files are used together as input for Model Selection.

We used SFS as input (x) in the NN, and the model names as the output (y) and removed the parameters as they are not necessary for this step. We used MinMax scaler from sklearn^75^ only on the SFS data as above, and the names of the models are converted to One-Hot Encoding by using pandas.Categorical and keras.utils.to_categorical. After concatenating files coming from all the competitive models, we randomized rows by a custom code^77^. We left around 10,000 random lines per each model to test the power of NN (as a testing dataset) and for ABC analysis and used the rest to train the NN (as a training dataset). The rest of the approach is exactly similar as before.

We used two hidden layers of the Dense neural network with the activation relu (Supplementary Figure 5). We used softmax for the output layer with the same number of units as the number of trained models. We added a Masking layer and a noise injection layer as above. We used a 1% dropout layer within every dense layer to make it more robust. We used categorical_crossentropy for the loss function from Keras and adam for the optimizer.

After the training was done, we used the testing dataset to predict models from the simulated SFS, which were then used to perform the cross-validation test using ABC with the tolerance of 0.0033 (which converts to 100 samples for three models) using mnlogistic. We calculated the model selection (abc.postpr) by using real data. See a schematic representation in Supplementary Figure 2. This approach is also similar to our previous study^19^.

#### Parameter Estimation with DLS

This method uses the Classic parameter estimation strategy of DL (described above) together with the SMC algorithm used for recursion. The approach here is close to the classic approach of SMC^47^ but not exactly the same.

We used the rejection method in ABC for parameter estimation as it generates posterior within the prior range with a tolerance of 0.05. We obtained the posterior range by taking the minimum and the maximum values from the ABC output. This range was then used as a prior range for the next iteration. This cycle repeated until shrinking for all parameters is more than 0.95, suggesting it has reached convergence.

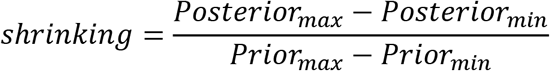

If the shrinking is more than 95% for a parameter, the new posterior estimation is rejected for that parameter. Instead, the prior is kept for another cycle. This strategy was used so that the posterior stops shrinking unless NN has found some accurate prediction for the parameter from SFS. We kept simulations inside the new posterior range in every cycle to reuse simulations from a previous cycle to a new cycle. A flow chart of this strategy can be found in Supplementary Figure 2c.

We used 10,000 simulations as a training dataset and 10,000 simulations for testing. The NN model is exactly as before (as used in DL, Supplementary Figure 5). To make it more efficient, we started with simulating a total length of 100 Mbp (each simulated region being 1 Mb long), and then we increased it stepwise (i.e., 0.5, 1.5, and 3 gbp). The priors for 100 Mbp regions are the same as presented in Table 1. The final posterior (after convergence reached) of 100 Mbp is used as a prior for 0.5 gbp simulation and so on. We multiplied the observed SFS by frac accordingly to scale it for to the simulated region length.

After the convergence was reached with 3 gbp in total, we finalized by running 50,000 training and 10,000 testing simulations with the DL method using loclinear with the tolerance of 0.01. The flowchart of the method is represented in Supplementary Figure 3.

#### Model Selection with DLS

Here we describe model selection using NN, ABC and SMC together. In principle, we can directly use the final output of parameter estimation by DLS for every model and then use it for the ABC classification approach. However, this approach would be inefficient, given that only one model is likely for our real dataset, and thus spending considerable resources to optimize parameters for unlikely scenarios does not make sense. Instead, we used the output of 100 Mbp parameter optimizations from the DLS approach as a prior to every model, and then we used the Model selection strategy of DL, as mentioned before. We found out that we already have enough power to distinguish between models using 100 Mbp for optimization.

### ABC-RF

We tested the real SFS against the three simulated models using a similar ABC approach but using Random Forest^78^ as an inferential tool implemented in the abcrf R package^48,79^. First, we trained our model using the bagging method applying the function abcrf, with no Linear Discriminant analysis, and 2,000 decision trees using 150,000 simulations (50,000 for each tested model). We then evaluated the performance of ABC-RF through a cross-validation dataset composed of 10,000 simulations for each tested model using the function predict.abcrf. The same function and settings were used for inferring the best-supported model using the SFS obtained from real data described above. We performed parameter estimation for the most supported scenario applying regression as implemented in the regAbcrf model using 1000 decision trees. Each parameter was inferred separately.

#### Relate

We used Relate^34^, a method for inferring local trees, to validate our parameter estimates. Relate uses branch length of the local trees to estimate coalescent rate through time^34^. Thus, we used it to compare effective population size (Ne) trajectories and inter-population coalescent rates for the African, European and East Asian populations between the real and simulated data. We applied Relate to YRI, CEU, and CHB samples (108, 99 and 103 individuals accordingly) from the high coverage version of the 1000 Genomes project as well as to genetic data simulated under each of the three models (Table 2, Supplementary Table 4 and 5). For real data chromosome 1 was used and a region of the same length was simulated.

We started with 2054 high-coverage genomes from the 1000 Genomes. We kept positions that a) are bi-allelic SNPs, b) pass the 1000 Genomes filter and have the QD (quality by depth) parameter above two and c) have a missing rate below 10%, We phased and imputed the entire dataset using Eagle version 2.4.1^80^. Next, we ran Relate on chromosome 1 for samples coming from the three focal populations. We used the GRCh38 recombination map, 1000 Genomes strict genomic mask and a mutation rate of 1.45×10^-8^. Next, we ran the Ne estimation module of Relate for each population individually for the Ne trajectory and for population pairs for the cross-coalescence curves.

For each model, we simulated a region of the same length as chromosome 1 with uniform recombination together for 100 African, 100 European and 100 East Asian individuals using msprime^18^. We used the 1000 Genomes strict mask for consistency between real and simulated data in terms of the length of the available sequence. After that, the simulated data were treated as described above.

We estimated Ne for both real and simulated data as 1/2C where C is the inferred intra-population coalescence rate. To estimate the relative inter-population coalescence rate, we used the following formula^33^:

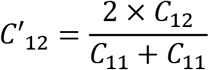

Where C_11_ and C_22_ are intra-population coalescence rates and C12 is the inter-population coalescence rate.

## Author Contributions

MM, FM and LP planned the project. MM, FM, VP, BY did the data analysis. MM, FM and VP wrote the paper. LP and BY corrected it.

## Acknowledgement

We kindly thank Michael Zody of New York Genome Center for giving us access to high coverage 1000 genome data. These data were generated at the New York Genome Center with funds provided by NHGRI Grant 3UM1HG008901-03S1. This research was supported by the European Union through Horizon 2020 research and innovation programme under grant no 810645, the European Regional Development Fund Project no. MOBEC008, 2014-2020.4.01.16-0030 and 2014-2020.4.01.16-0024.

## Code availability

We implemented a python based software package in https://github.com/mayukhmondal/ABC-DLS. Although it is python based the entire package can be accessible through the command line.

## Competing interests

The authors declare no competing interests.

## Notes

### Competing Interest Statement

The authors have declared no competing interest.

### Summary of Updates

Update with correct Figure 1

## Reference

1. Hublin, J.-J. et al. New fossils from Jebel Irhoud, Morocco and the pan-African origin of Homo sapiens. Nature 546, 289–292 (2017).

2. Trinkaus, E. Femoral neck-shaft angles of the Qafzeh-Skhul early modern humans, and activity levels among immature Near Eastern Middle Paleolithic hominids. J. Hum. Evol. 25, 393–416 (1993).

3. Liu, W. et al. The earliest unequivocally modern humans in southern China. Nature 526, 696–699 (2015).

4. Harvati, K. et al. Apidima Cave fossils provide earliest evidence of Homo sapiens in Eurasia. Nature 571, 500–504 (2019).

5. Mondal, M. et al. Genomic analysis of Andamanese provides insights into ancient human migration into Asia and adaptation. Nat. Genet. 48, 1066–70 (2016).

6. Malaspinas, A. S. et al. A genomic history of Aboriginal Australia. Nature 538, 1–152 (2016).

7. Mallick, S. et al. The Simons Genome Diversity Project: 300 genomes from 142 diverse populations. Nature (2016) doi:10.1038/nature18964.

8. Clarkson, C. et al. Human occupation of northern Australia by 65,000 years ago. Nature 547, 306–310 (2017).

9. Groucutt, H. S. et al. Homo sapiens in Arabia by 85,000 years ago. Nat. Ecol. Evol. 2, 800–809 (2018).

10. Cabrera, V. M., Marrero, P., Abu-Amero, K. K. & Larruga, J. M. Carriers of mitochondrial DNA macrohaplogroup L3 basal lineages migrated back to Africa from Asia around 70,000 years ago. BMC Evol. Biol. 18, 98 (2018).

11. Poznik, G. D. et al. Punctuated bursts in human male demography inferred from 1,244 worldwide Y-chromosome sequences. Nat. Genet. 48, 593–599 (2016).

12. Ciregan, D., Meier, U. & Schmidhuber, J. Multi-column deep neural networks for image classification. in 2012 IEEE conference on computer vision and pattern recognition 3642–3649 (IEEE, 2012).

13. Graves, A. & Schmidhuber, J. Offline handwriting recognition with multidimensional recurrent neural networks. in Advances in neural information processing systems 545–552 (2009).

14. Hutter, M. On universal prediction and Bayesian confirmation. Theor. Comput. Sci. 384, 33–48 (2007).

15. Kurtz, D. M. et al. Dynamic risk profiling using serial tumor biomarkers for personalized outcome prediction. Cell 178, 699–713 (2019).

16. Goldberg, Y. A primer on neural network models for natural language processing. J. Artif. Intell. Res. 57, 345–420 (2016).

17. Hudson, R. R. Generating samples under a Wright-Fisher neutral model of genetic variation. Bioinforma. Oxf. Engl. 18, 337–338 (2002).

18. Kelleher, J., Etheridge, A. M. & McVean, G. Efficient Coalescent Simulation and Genealogical Analysis for Large Sample Sizes. PLoS Comput. Biol. 12, 1–22 (2016).

19. Mondal, M., Bertranpetit, J. & Lao, O. Approximate Bayesian computation with deep learning supports a third archaic introgression in Asia and Oceania. Nat. Commun. 10, (2019).

20. Jay, F., Boitard, S. & Austerlitz, F. An ABC method for whole-genome sequence data: Inferring Paleolithic and Neolithic human expansions. Mol. Biol. Evol. 36, 1565–1579 (2019).

21. Villanea, F. A. & Schraiber, J. G. Multiple episodes of interbreeding between Neanderthal and modern humans. Nat. Ecol. Evol. 3, 39–44 (2019).

22. Kern, A. D. & Schrider, D. R. diploS/HIC: an updated approach to classifying selective sweeps. G3 Genes Genomes Genet. 8, 1959–1970 (2018).

23. Torada, L. et al. ImaGene: a convolutional neural network to quantify natural selection from genomic data. BMC Bioinformatics 20, 337 (2019).

24. Sanchez, T., Cury, J., Charpiat, G. & Jay, F. Deep learning for population size history inference: design, comparison and combination with approximate Bayesian computation. bioRxiv 2020.01.20.910539 (2020) doi:10.1101/2020.01.20.910539.

25. Schlebusch, C. M. et al. Southern African ancient genomes estimate modern human divergence to 350,000 to 260,000 years ago. Science 358, 652–655 (2017).

26. Grün, R. et al. U-series and ESR analyses of bones and teeth relating to the human burials from Skhul. J. Hum. Evol. 49, 316–334 (2005).

27. Pagani, L. et al. Genomic analyses inform on migration events during the peopling of Eurasia. Nature 538, 238–242 (2016).

28. Kuhlwilm, M. et al. Ancient gene flow from early modern humans into Eastern Neanderthals. Nature (2016) doi:10.1038/nature16544.

29. Soares, P. et al. The expansion of mtDNA haplogroup L3 within and out of Africa. Mol. Biol. Evol. 29, 915–927 (2012).

30. Karmin, M. et al. A recent bottleneck of Y chromosome diversity coincides with a global change in culture. Genome Res. 25, 459–466 (2015).

31. Mondal, M. et al. Y-chromosomal sequences of diverse Indian populations and the ancestry of the Andamanese. Hum. Genet. 136, (2017).

32. Haber, M. et al. A rare deep-rooting D0 African Y-chromosomal haplogroup and its implications for the expansion of modern humans out of Africa. Genetics 212, 1421–1428 (2019).

33. Schiffels, S. & Durbin, R. Inferring human population size and separation history from multiple genome sequences. Nat. Genet. 46, 919–925 (2014).

34. Speidel, L., Forest, M., Shi, S. & Myers, S. R. A method for genome-wide genealogy estimation for thousands of samples. Nat. Genet. 51, 1321–1329 (2019).

35. Cole, C. B., Zhu, S. J., Mathieson, I., Prüfer, K. & Lunter, G. Ancient Admixture into Africa from the ancestors of non-Africans. bioRxiv (2020) doi:10.1101/2020.06.01.127555.

36. Lipson, M. et al. Ancient West African foragers in the context of African population history. Nature 577, 665–670 (2020).

37. Gutenkunst, R. N., Hernandez, R. D., Williamson, S. H. & Bustamante, C. D. Inferring the joint demographic history of multiple populations from multidimensional SNP frequency data. PLoS Genet. 5, (2009).

38. Li, H. & Durbin, R. Inference of Human Population History From Whole Genome Sequence of A Single Individual. Nature 475, 493–496 (2012).

39. Gravel, S. et al. Demographic history and rare allele sharing among human populations. Proc. Natl. Acad. Sci. U. S. A. 108, 11983–11988 (2011).

40. Information, S. Geographical barriers, environmental challenges, and complex migration events during the peopling of Eurasia.

41. Schlebusch, C. M. et al. Ancient genomes from southern Africa pushes modern human divergence beyond 260,000 years ago. Doi.Org 655, 145409 (2017).

42. Loosdrecht, M. van de et al. Pleistocene North African genomes link Near Eastern and sub-Saharan African human populations. Science 360, 548–552 (2018).

43. Collette, A. Python and HDF5: unlocking scientific data. (O’Reilly Media, Inc., 2013).

44. Abadi, M. et al. TensorFlow: A system for large-scale machine learning. in Proceedings of the 12th USENIX Symposium on Operating Systems Design and Implementation, OSDI 2016 (2016).

45. Chollet, F. and others. Home - Keras Documentation. https://keras.io/ (2015).

46. Sisson, S. A., Fan, Y. & Tanaka, M. M. Sequential monte carlo without likelihoods. Proc. Natl. Acad. Sci. 104, 1760–1765 (2007).

47. Liu, J. S. & Chen, R. Sequential Monte Carlo methods for dynamic systems. J. Am. Stat. Assoc. 93, 1032–1044 (1998).

48. Raynal, L. et al. ABC random forests for Bayesian parameter inference. Bioinformatics 35, 1720–1728 (2019).

49. Jouganous, J., Long, W. & Gravel, S. Inferring the Joint Demographic History of Multiple Populations: Beyond the Diffusion Approximation. 1–37 (2017).

50. Green, R. E. et al. A draft sequence of the Neandertal genome. Science 328, 710–722 (2010).

51. Browning, S. R., Browning, B. L., Zhou, Y., Tucci, S. & Akey, J. M. Analysis of Human Sequence Data Reveals Two Pulses of Archaic Denisovan Admixture. Cell 173, 53–61.e9 (2018).

52. Jacobs, G. S. et al. Multiple Deeply Divergent Denisovan Ancestries in Papuans. Cell 177, 1010–1021.e32 (2019).

53. Ragsdale, A. P. & Gravel, S. Models of archaic admixture and recent history from two-locus statistics. PLoS Genet. 15, 1–19 (2019).

54. Lorente-galdos, B. et al. Whole-genome sequence analysis of a Pan African set of samples reveals archaic gene flow from an extinct basal population of modern humans into sub-Saharan populations. 1–15 (2019).

55. Chen, L., Wolf, A. B., Fu, W., Li, L. & Akey, J. M. Identifying and Interpreting Apparent Neanderthal Ancestry in African Individuals. Cell 180, 677–687.e16 (2020).

56. Bergström, A. et al. Insights into human genetic variation and population history from 929 diverse genomes. Science 367, (2020).

57. Scally, A. The mutation rate in human evolution and demographic inference. Curr. Opin. Genet. Dev. 41, 36–43 (2016).

58. Kong, A. et al. Rate of de novo mutations and the importance of father’s age to disease risk. Nature 488, 471–475 (2012).

59. Tian, X., Browning, B. L. & Browning, S. R. Estimating the genome-wide mutation rate with three-way identity by descent. Am. J. Hum. Genet. 105, 883–893 (2019).

60. Excoffier, L., Dupanloup, I., Huerta-Sánchez, E., Sousa, V. C. & Foll, M. Robust Demographic Inference from Genomic and SNP Data. PLoS Genet. 9, (2013).

61. Theunert, C., Tang, K., Lachmann, M., Hu, S. & Stoneking, M. Inferring the History of Population Size Change from Genome-Wide SNP Data Research article. 29, 3653–3667 (2012).

62. Wall, J. D. Inferring Human Demographic Histories of Non-African Populations from Patterns of Allele Sharing. Am. J. Hum. Genet. (2017) doi:10.1016/j.ajhg.2017.04.002.

63. Lapierre, M., Lambert, A. & Achaz, G. Accuracy of demographic inferences from the site frequency spectrum: The case of the yoruba population. Genetics 206, 139–449 (2017).

64. Meyer, M. et al. A High-Coverage Genome Sequence from an Archaic Denisovan Individual. Science 338, 222–226 (2012).

65. Mondal, M., Casals, F., Majumder, P. P. & Bertranpetit, J. Reply to ‘No evidence for unknown archaic ancestry in South Asia’. Nat. Genet. 50, 1637–1639 (2018).

66. Moorjani, P. Molecular clock helps estimate age of ancient genomes. PNaS 113, 5459–5460 (2016).

67. Besenbacher, S., Hvilsom, C., Marques-Bonet, T., Mailund, T. & Schierup, M. H. Direct estimation of mutations in great apes reconciles phylogenetic dating. Nat. Ecol. Evol. 3, 286–292 (2019).

68. Pagani, L. & Crevecoeur, I. What is Africa? A human perspective. (2018). doi:10.13140/RG.2.2.16605.10728.

69. Tremblay, M. & Vézina, H. New Estimates of Intergenerational Time Intervals for the Calculation of Age and Origins of Mutations. Am. J. Hum. Genet. 66, 651–658 (2000).

70. Auton, A. et al. A global reference for human genetic variation. Nature 526, 68–74 (2015).

71. Li, H. A statistical framework for SNP calling, mutation discovery, association mapping and population genetical parameter estimation from sequencing data. Bioinformatics 27, 2987–2993 (2011).

72. Danecek, P. et al. The variant call format and VCFtools. Bioinformatics 27, 2156–2158 (2011).

73. Alistair Miles et al. cggh/scikit-allel: v1.3.2. (Zenodo, 2020). doi:10.5281/zenodo.3976233.

74. Köster, J. & Rahmann, S. Snakemake—a scalable bioinformatics workflow engine. Bioinformatics 28, 2520–2522 (2012).

75. Pedregosa, F. et al. Scikit-learn: Machine learning in Python. J. Mach. Learn. Res. 12, 2825–2830 (2011).

76. Csilléry, K., François, O. & Blum, M. Approximate Bayesian Computation (ABC) in R: A Vignette. 202.162.217.53 1–21 (2012).

77. Salle, A., Idiart, M. & Villavicencio, A. Matrix factorization using window sampling and negative sampling for improved word representations. ArXiv Prepr. ArXiv160600819 (2016).

78. Breiman, L. Random forests. Mach. Learn. 45, 5–32 (2001).

79. Pudlo, P. et al. Reliable ABC model choice via random forests. Bioinformatics 32, 859–866 (2016).

80. Loh, P.-R., Palamara, P. F. & Price, A. L. Fast and accurate long-range phasing in a UK Biobank cohort. Nat. Genet. 48, 811–816 (2016).

