## Supplementary Information for "Revisiting the Out of Africa event with a novel Deep Learning approach"

* Contributed Equally

Correspondence and requests for materials should be addressed MM (email: mondal [dot] mayukh [at] gmail [dot] com).

### Supplementary Figures

#### Supplementary Figure 1: The simplistic schema of the models

a) Simple Out of Africa (model S), b) Back to Africa (model B) and c) Out of Africa Mixed (model M)

a)

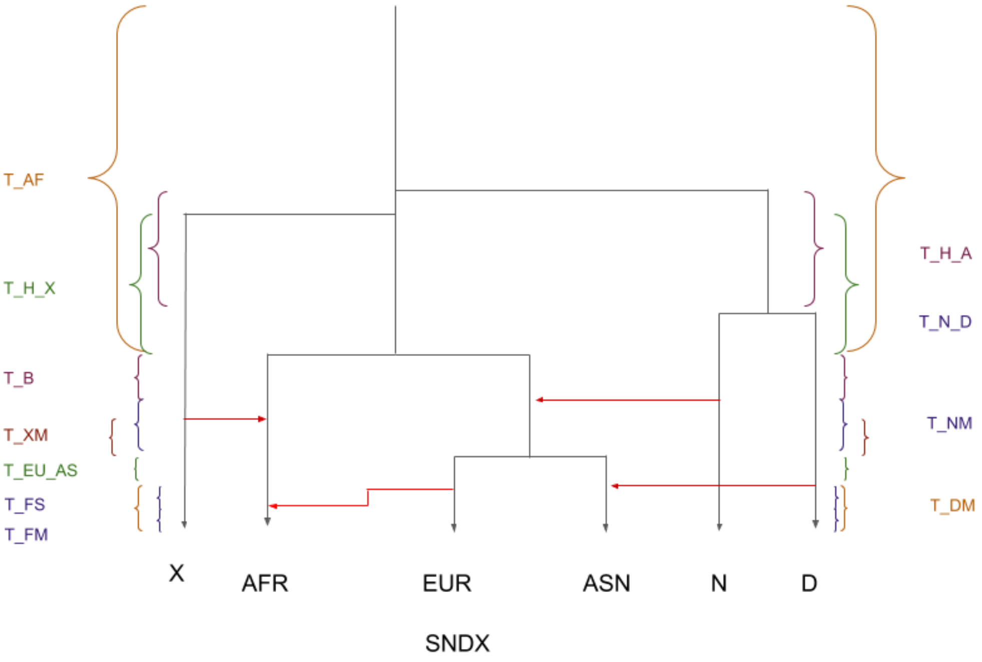


b)

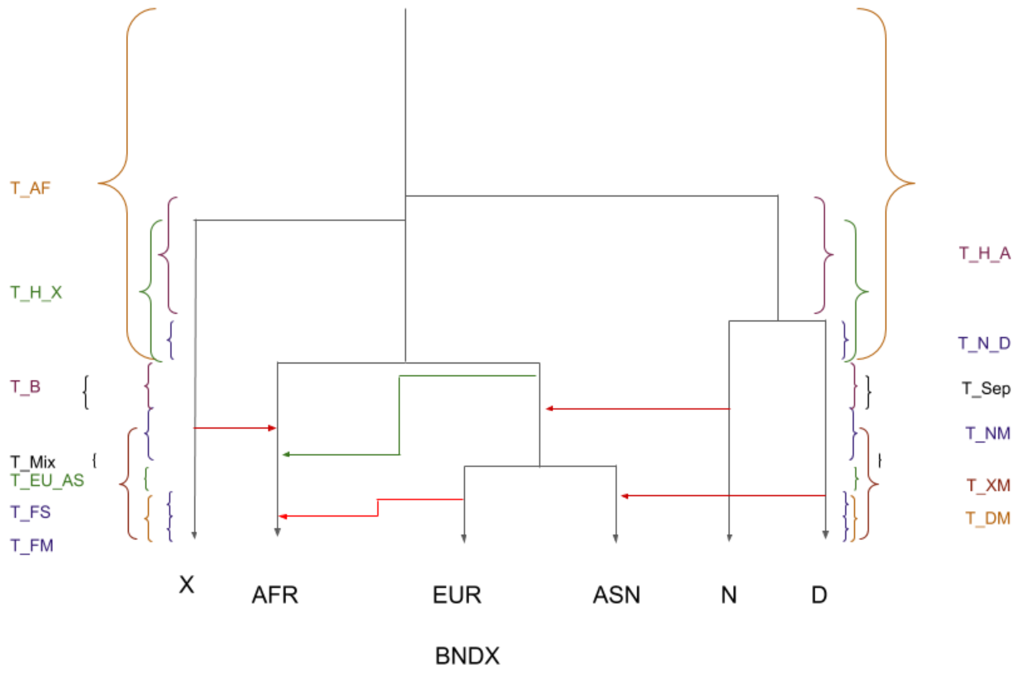
T_AF
T_H_X
T_B
T_Mix
T_EU_AS
T_FS
T_FM
T_H_A
T_N_D
T_Sep
T_NM
T_XM
T_DM
X
AFR
EUR
ASN
N
D
BNDX

c)

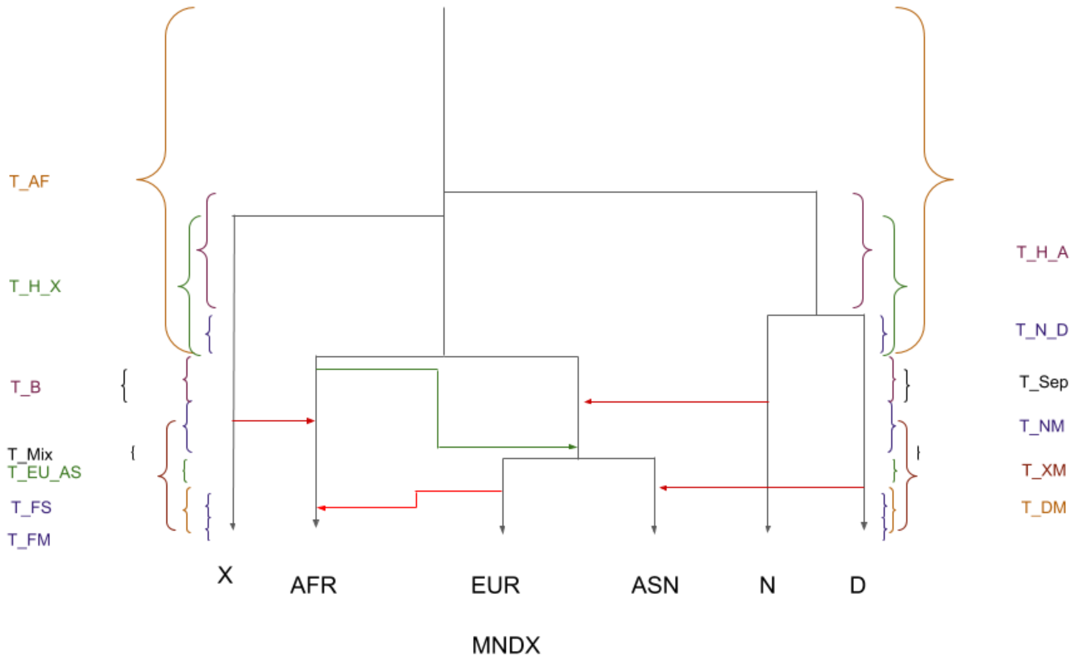
T_AF
T_H_X
T_B
T_Mix
T_EU_AS
T_FS
T_FM
T_H_A
T_N_D
T_Sep
T_NM
T_XM
T_DM
X
AFR
EUR
ASN
N
D
MNDX

#### Supplementary Figure 2: Flowchart of individual ABC-DLS method

a) Model Selection by DLS, b) Param Prediction by DL c) SMC

a)

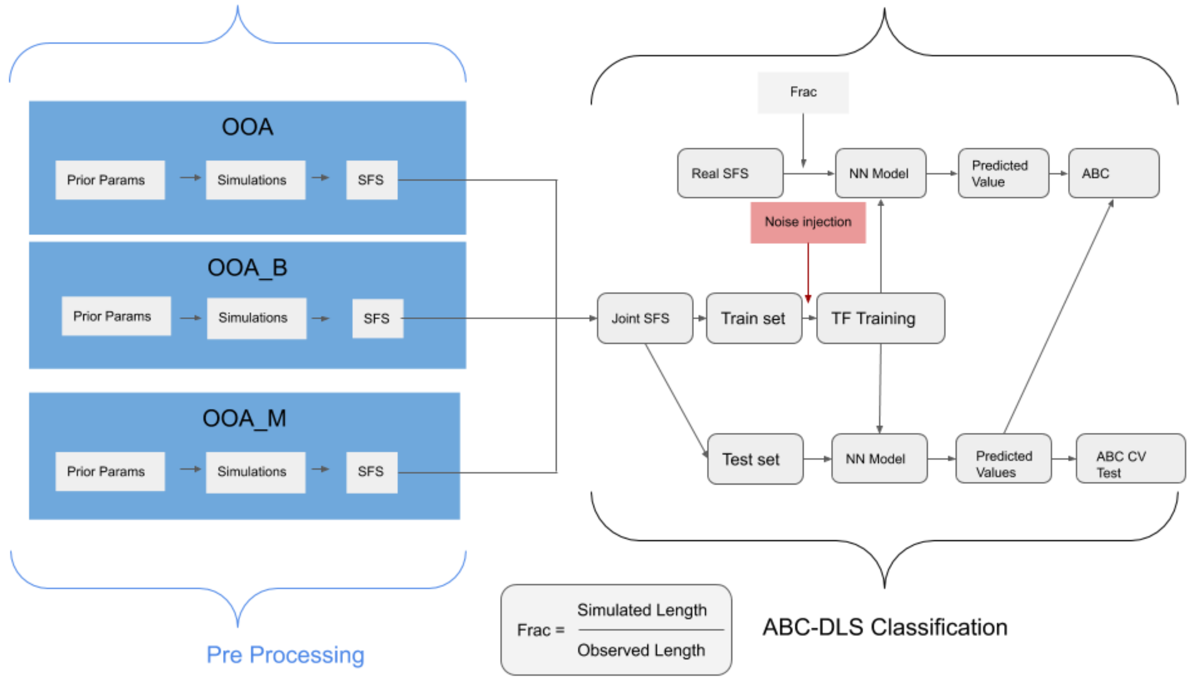


b)

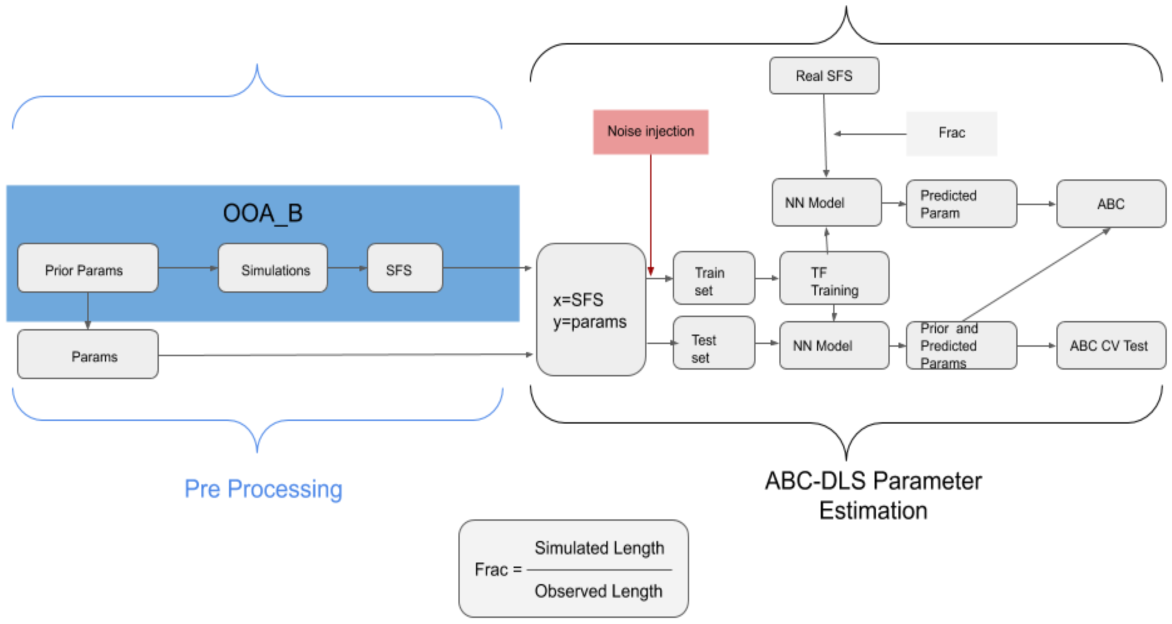

c)

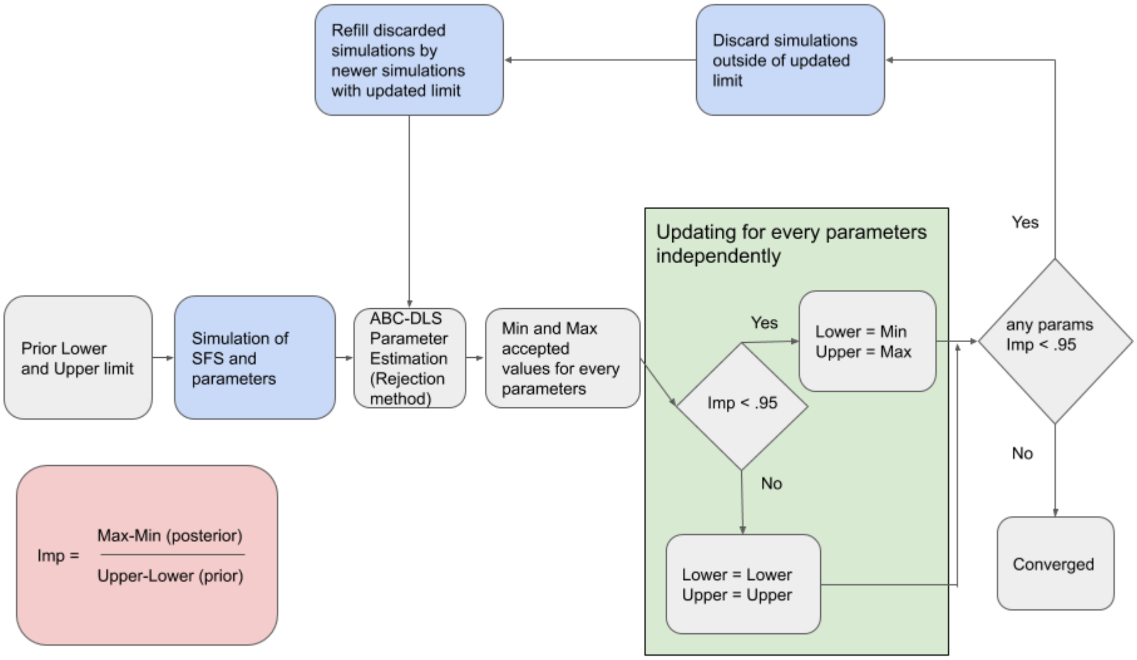

Refill discarded simulations by newer simulations with updated limit
Discard simulations outside of updated limit
Updating for every parameters independently
Yes
Prior Lower and Upper limit
Simulation of SFS and parameters
ABC-DLS Parameter Estimation (Rejection method)
Min and Max accepted values for every parameters
Yes
Lower = Min
Upper = Max
any params Imp < .95
Imp < .95
No
No
Lower = Lower
Upper = Upper
Converged
Imp = Max-Min (posterior) / Upper-Lower (prior)

#### Supplementary Figure 3: Flowchart for the ABC-DLS methods interacting together to get desired results

a) Parameter Estimation using DLS b) Model Selection using DLS

a)

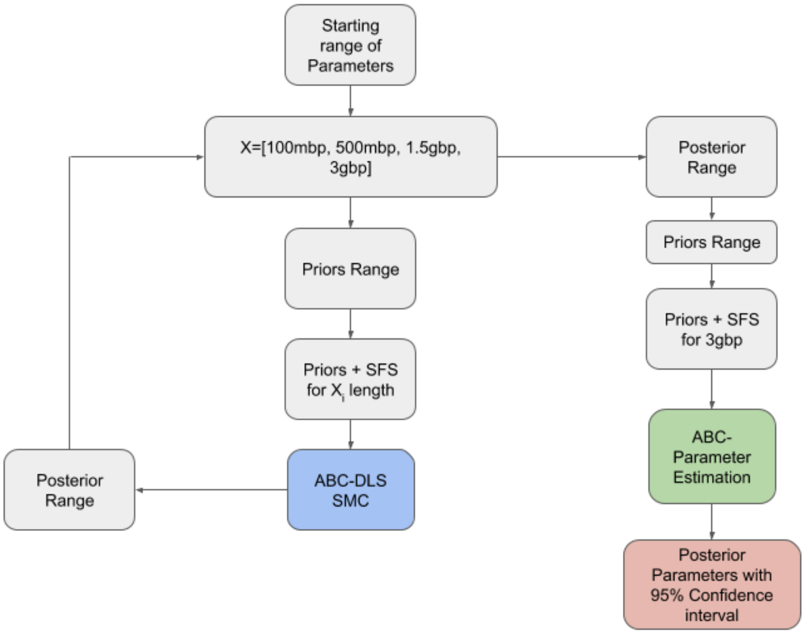


b)

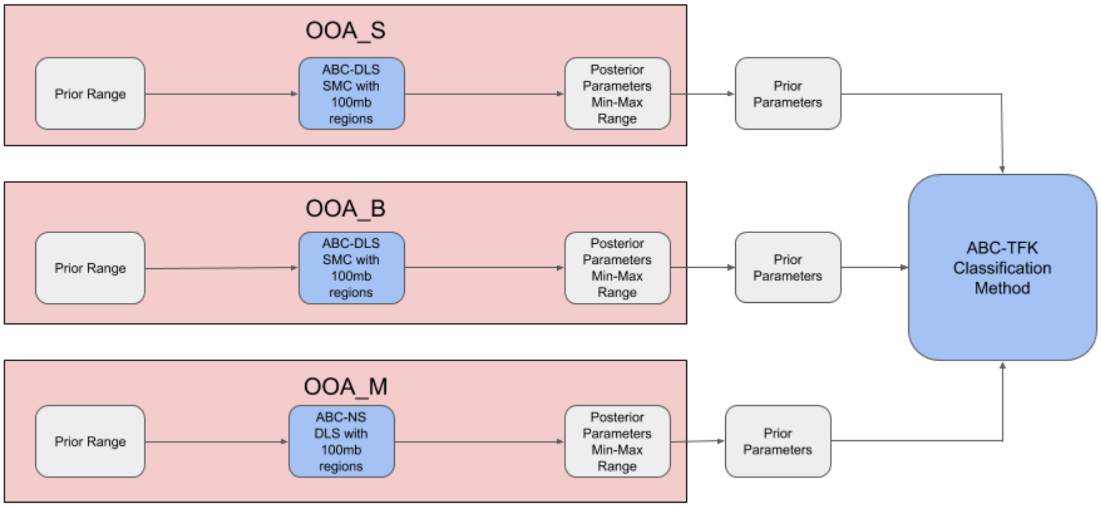

Supplementary Figure 4: Cross population coalescent rate calculated using relate for Model S

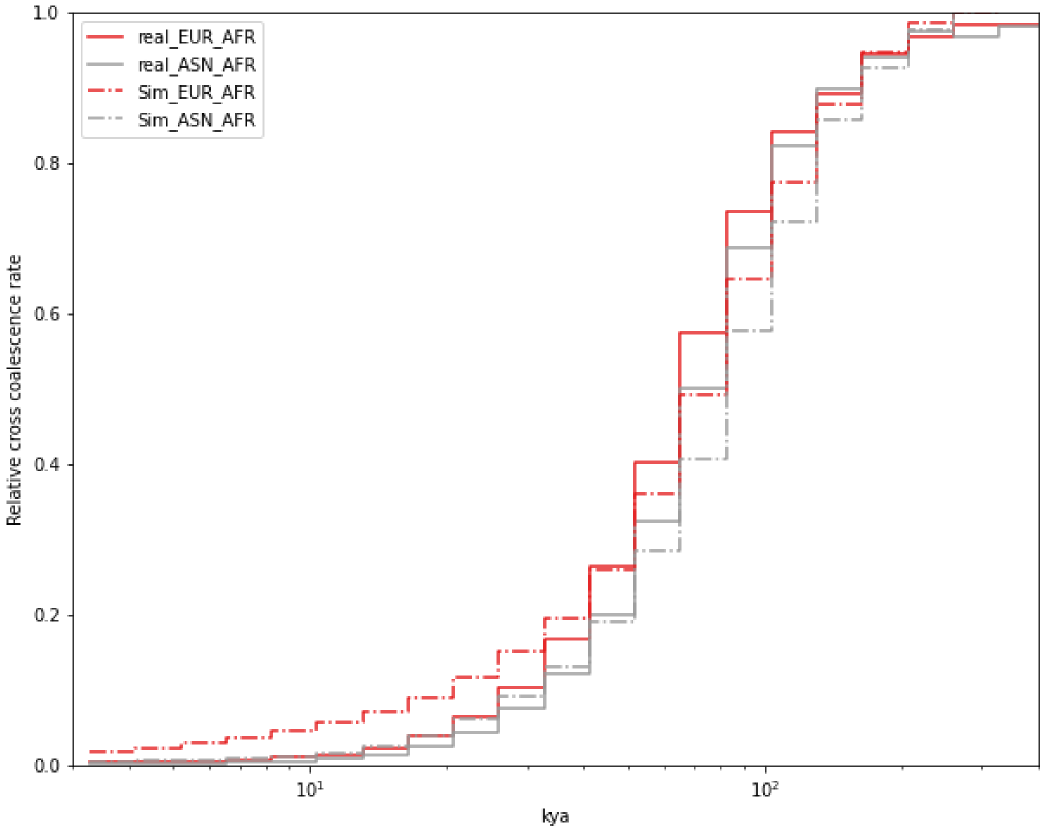

#### Supplementary Figure 5: Schematic for used TensorFlow model

a) Model Selection and b) Parameter Estimation. "?" marks the number of rows which can be variable depending on the simulation and input files. The number denotes the number of elements for input and output per simulation.

a)

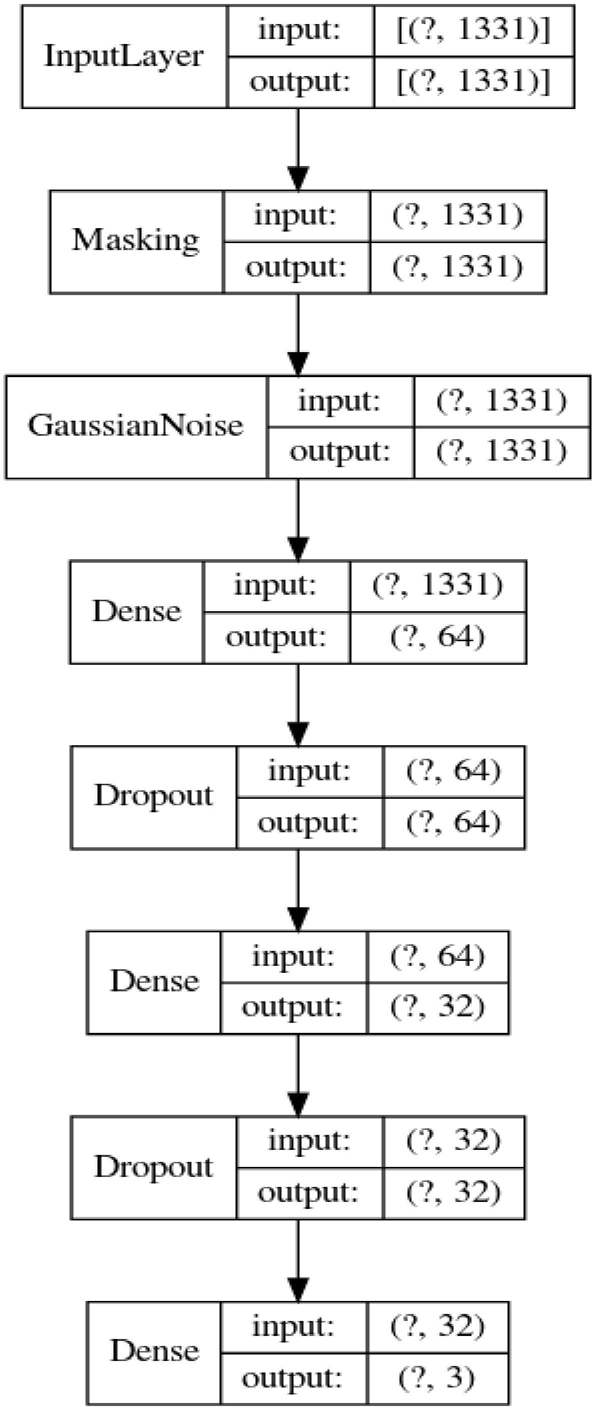

b)

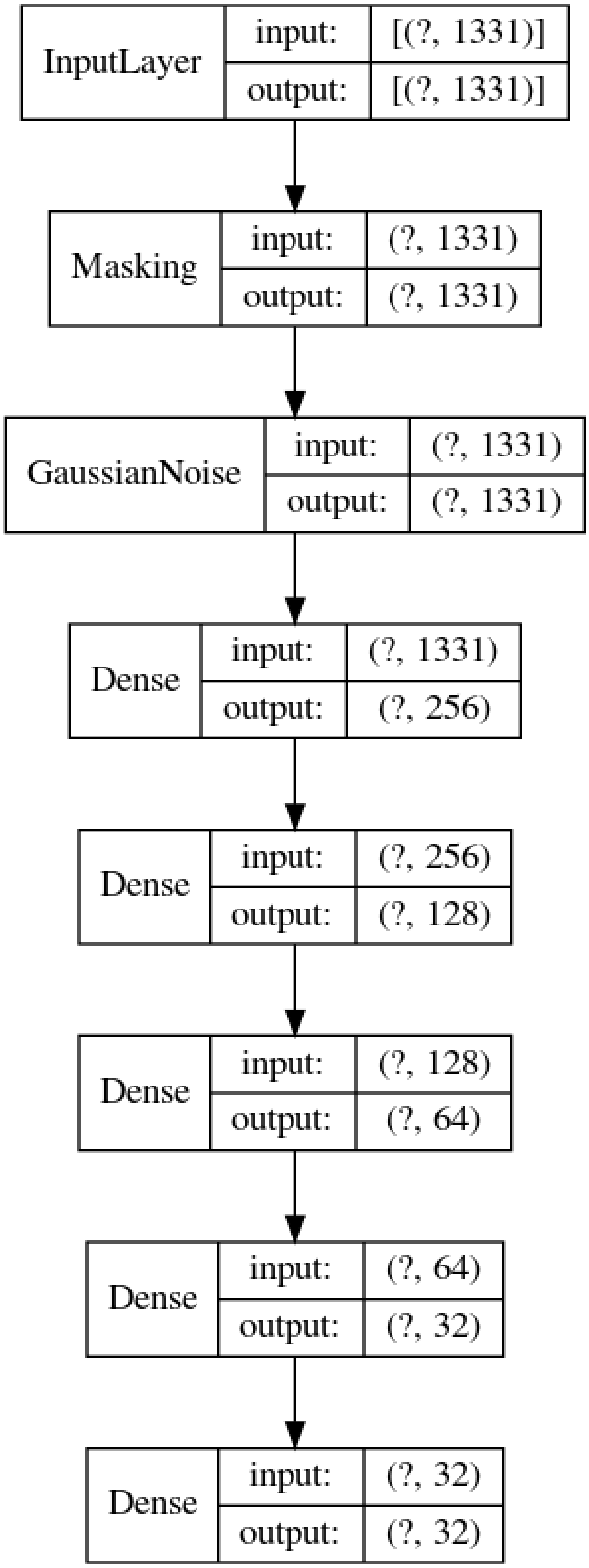

InputLayer
input:
[(?, 1331)]
output:
[(?, 1331)]
Masking
input:
(?, 1331)
output:
(?, 1331)
GaussianNoise
input:
(?, 1331)
output:
(?, 1331)
Dense
input:
(?, 1331)
output:
(?, 256)
Dense
input:
(?, 256)
output:
(?, 128)
Dense
input:
(?, 128)
output:
(?, 64)
Dense
input:
(?, 64)
output:
(?, 32)
Dense
input:
(?, 32)
output:
(?, 32)

### Supplementary Tables

Supplementary Table 1: Prior and Posterior range of Out Of Africa model with migrations.

| Parameters | Prior | Posterior |
|---|---|---|
| **N_A** | 5000 - 25000 | 15400 (15372 - 15413) |
| **N_AF** | 10000 - 150000 | 21321 (21241 - 21391) |
| **N_EU** | 10000 - 150000 | 96561 (92254 - 100026) |
| **N_AS** | 10000 - 150000 | 95454 (92322 - 98768) |
| **N_EU0** | 500 - 5000 | 1021 (1007 - 1031) |
| **N_AS0** | 500 - 5000 | 622 (615 - 628) |
| **N_B** | 500 - 5000 | 3629 (3623 - 3643) |
| **T_EU_AS (ky)** | 15 - 80 | 41.37 (41.21 - 41.53) |
| **T_B (ky)** | 5 - 320 | 315.97 (300.19 - 321.87) |
| **T_AF (ky)** | 5 - 700 | 252.78 (247.79 - 263.28) |
| **m_AF_B ($\times 10^{-5}$)** | 0 - 50 | 21.66 (21.5 - 21.75) |
| **m_AF_EU ($\times 10^{-5}$)** | 0 - 50 | 1.3 (1.22 - 1.35) |
| **m_AF_AS ($\times 10^{-5}$)** | 0 - 50 | 0.3 (0.27 - 0.37) |
| **m_EU_AS ($\times 10^{-5}$)** | 0 - 50 | 10.29 (10.11 - 10.44) |

Posterior column represents Mean and CI (Confidence Interval of 2.5%-97.5%) of respective parameters.

#### Supplementary Table 2: Posterior range of model S.

| **Parameters** | **Mean** | **CI** | **Events (kya)** |
|---|---|---|---|
| **N_A** | 13,921 | 13,917 - 13,922 | |
| **N_AF** | 16,064 | 15,984 - 16,142 | |
| **N_EU** | 112,893 | 106,050 - 121,497 | |
| **N_AS** | 134,030 | 127,105 - 142,323 | |
| **N_F** | 17,099 | 5,152 - 29,627 | |
| **N_EU0** | 1,844 | 1,814 - 1,898 | |
| **N_AS0** | 760 | 750 - 779 | |
| **N_B** | 1,356 | 1,322 - 1,383 | |
| **T_FM (ky)** | 4 | 2.4 - 4.9 | 4 (2.4 - 4.9) |
| **T_FS (ky)** | 6 | 0.6 - 9.9 | 9.5 (4.7 - 14.3) |
| **T_DM (ky)** | 15.4 | 15.2 - 15.6 | 15.4 (15.2 - 15.6) |
| **T_EU_AS (ky)** | 18.1 | 18 - 18.4 | 33.5 (33.2 - 33.8) |
| **T_XM (ky)** | 5.2 | 5.1 - 5.5 | 38.7 (38.3 - 39.1) |
| **T_NM (ky)** | 6.4 | 6 - 6.8 | 39.9 (39.4 - 40.4) |
| **T_B (ky)** | 10.9 | 10.5 - 11.5 | 50.8 (50.1 - 51.6) |
| **T_AF (ky)** | 327.3 | 316.1 - 335 | 378.1 (368.7 - 387.5) |
| **T_N_D (ky)** | 454 | 453.8 - 454.2 | 454 (453.8 - 454.2) |
| **T_H_A (ky)** | 253.1 | 252.8 - 253.2 | 707.1 (706.8 - 707.4) |
| **T_H_X (ky)** | 684.1 | 677.1 - 687.8 | 684.1 (677.1 - 687.8) |
| **NMix (%)** | 3.03 | 3.03 - 3.03 | |
| **DMix (%)** | 0.49 | 0.45 - 0.51 | |
| **XMix (%)** | 6.05 | 5.88 - 6.14 | |
| **FMix (%)** | 6.28 | 6.09 - 6.47 | |

CI is the confidence interval of 2.5%-97.5% of respective parameters. Ky is kilo years and kya is kilo years ago from now.

#### Supplementary Table 3: Posterior range of model M.

| Parameters | Mean | CI | Events (kya) |
|---|---|---|---|
| N_A | 13,706 | 13,676 - 13,744 | |
| N_AF | 16,163 | 16,037 - 16,319 | |
| N_EU | 95,459 | 80,211 - 109,645 | |
| N_AS | 96,852 | 85,077 - 110,425 | |
| N_F | 17,949 | 5,624 - 29,178 | |
| N_EU0 | 1,716 | 1,645 - 1,799 | |
| N_AS0 | 673 | 651 - 696 | |
| N_MX | 519 | 509 - 533 | |
| N_B | 4,486 | 4,013 - 4,836 | |
| N_B0 | 16,541 | 8,963 - 22,766 | |
| T_FM (ky) | 3.6 | 2.0 - 5 | 3.6 (2 - 5) |
| T_FS (ky) | 5.5 | 0.8 - 9.7 | 9.1 (4.5 - 13.8) |
| T_DM (ky) | 12.2 | 10.8 - 13.9 | 12.2 (10.8 - 13.9) |
| T_EU_AS (ky) | 16.2 | 14.8 - 17.7 | 28.4 (26.3 - 30.5) |
| T_NM (ky) | 5.3 | 5.1 - 5.8 | 33.74 (31.6 - 35.9) |
| T_XM (ky) | 27.2 | 26.3 - 29 | 55.6 (53.1 - 58.1) |
| T_Mix (ky) | 5.6 | 5.3 - 5.9 | 34.1 (31.9 - 36.2) |
| T_Sep (ky) | 9.2 | 9.0 - 9.4 | 64.8 (62.3 - 67.3) |
| T_B (ky) | 8.4 | 5.7 - 13.7 | 73.2 (68.5 - 77.9) |
| T_AF (ky) | 79.4 | 73.9 - 94.4 | 152.6 (141.4 - 163.9) |
| T_N_D (ky) | 381.2 | 334.1 - 428.8 | 381.2 (334.1 - 428.8) |
| T_H_A (ky) | 179.7 | 135.1 - 240.1 | 560.9 (490.2 - 631.6) |
| T_H_X (ky) | 692.2 | 683.6 - 698.3 | 692.2 (683.6 - 698.3) |
| Mix (%) | 88.29% | 87.75 - 89.2 | |
| NMix (%) | 1.67% | 1.06 - 2.24 | |
| DMix (%) | 0.63% | 0.56 - 0.73 | |
| XMix (%) | 8.59% | 8.47 - 8.76 | |
| FMix (%) | 5.32% | 5.15 - 5.51 | |

CI is the confidence interval of 2.5%-97.5% of respective parameters. Ky is kilo years and kya is kilo years ago from now.

Supplementary Table 4: Posterior range of model B with migration rates with known parameters coming from a simulation from Table 1.

| Parameters | Mean | 2.5P | 97.5P |
|---|---|---|---|
| N_A | 13,278 | 13172 | 13,403 |
| N_AF | 16,080 | 12696 | 20,214 |
| N_EU | 83,243 | 25183 | 150,157 |
| N_AS | 69,722 | 42271 | 109,262 |
| N_EU0 | 1,589 | 1128 | 2,234 |
| N_AS0 | 646 | 552 | 764 |
| N_BC | 16,659 | 4176 | 30,054 |
| N_B | 2,665 | 2406.00 | 2,929 |
| N_AF0 | 24,791 | 20477.00 | 29,208 |
| T_DM (ky) | 19.2 | 16.6 | 22.6 |
| T_EU_AS (ky) | 10.6 | 8.1 | 13.5 |
| T_NM (ky) | 7.5 | 5.4 | 9.7 |
| T_XM (ky) | 17.8 | 10.8 | 27.3 |
| T_Mix (ky) | 21.6 | 14.6 | 28.7 |
| T_Sep (ky) | 12.3 | 5.9 | 19.2 |
| T_B (ky) | 17.9 | 12.8 | 24.1 |
| T_AF (ky) | 251.8 | 208.0 | 294.8 |
| T_N_D (ky) | 428.0 | 401.6 | 448.7 |
| T_H_A (ky) | 227.9 | 200.5 | 248.2 |
| T_H_X (ky) | 599.6 | 524.9 | 691.8 |
| m_AF_B (×10-5) | 12 | 1.95 | 22.2 |
| m_B_AF (×10-5) | 4 | 1.91 | 6.27 |
| m_AF_EU (×10-5) | 2 | 0.28 | 3.1 |
| m_EU_AF (×10-5) | 0 | 0.14 | 0.88 |
| m_AF_AS (×10-5) | 1.41 | 0.31 | 2.38 |
| m_AS_AF (×10-5) | 0.39 | 0.08 | 0.72 |
| m_EU_AS (×10-5) | 4.92 | 2.11 | 7.84 |
| m_AS_EU (×10-5) | 2.11 | 0.35 | 3.95 |
| Mix (%) | 79.91% | 64.24% | 92.69% |
| NMix (%) | 2.69% | 2.36% | 2.97% |
| DMix (%) | 1.11% | 0.87% | 1.32% |
| XMix (%) | 6.60% | 4.05% | 9.67% |

2.5P is 2.5 percentile and 97.5P is 97.5 percentile. Ky is kilo years.

Supplementary Table 5: Posterior range of model S with migration rates with known parameters coming from a simulation from Table 1.

| Parameters | Mean | 2.5P | 97.5P |
|---|---|---|---|
| **N_A** | 13,390 | 12,895 | 13,874 |
| **N_AF** | 15,137 | 14,833 | 15,381 |
| **N_EU** | 95,604 | 55,916 | 149,202 |
| **N_AS** | 83,758 | 59,848 | 108,935 |
| **N_EU0** | 1,593 | 1,319 | 1,900 |
| **N_AS0** | 696 | 642 | 768 |
| **N_B** | 1853 | 1,783 | 1,952 |
| **T_DIntro** | 16.1 | 13.9 | 18.3 |
| **T_EU_AS** | 14.7 | 12.3 | 17.3 |
| **T_NIntro** | 5.8 | 5.2 | 6.6 |
| **T_XIntro** | 10.2 | 6.1 | 14.3 |
| **T_B** | 18.8 | 14.8 | 23.7 |
| **T_AF** | 498.1 | 319.6 | 684.2 |
| **T_N_D** | 429.6 | 399.4 | 449.8 |
| **T_H_A** | 229.9 | 203.4 | 249.0 |
| **T_H_X** | 669.4 | 633.4 | 697.0 |
| **m_AF_B** | 6.906 | 6.5658 | 7.1199 |
| **m_B_AF** | 10.66 | 9.86 | 11.59 |
| **m_AF_EU** | 6.40 | 5.96 | 6.92 |
| **m_EU_AF** | 0.28 | 0.05 | 0.53 |
| **m_AF_AS** | 1.56 | 1.15 | 2.07 |
| **m_AS_AF** | 0.19 | 0.05 | 0.33 |
| **m_EU_AS** | 2.71 | 1.14 | 4.13 |
| **m_AS_EU** | 1.40 | 0.33 | 2.60 |
| **nintro** | 2.38% | 2.08% | 2.70% |
| **dintro** | 0.78% | 0.68% | 0.83% |
| **xintro** | 4.68% | 4.18% | 5.09% |

2.5P is 2.5 percentile and 97.5P is 97.5 percentile. Ky is kilo years.

Supplementary Table 6: Cross validation and Model Selection under different strategy for real data.

| **Filtering Strategy** | B | M | S |
|---|---|---|---|
| B | 99.13% | 0.00% | 0.87% |
| M | 0.00% | 100.00% | 0.00% |
| S | 0.09% | 0.02% | 99.89% |
| Posterior model probabilities | 100.00% | 0.00% | 0.00% |
| **Dataset** | | | |
| B | 100.00% | 0.00% | 0.00% |
| M | 0.00% | 100.00% | 0.00% |
| S | 0.67% | 0.06% | 99.27% |
| Posterior model probabilities | 100.00% | 0.00% | 0.00% |

Confusion matrix for misclassification is reported here using SMC for random samples from the models. Posterior model probabilities are final posterior after using the real data.

Supplementary Table 7: Cross validation and Model Selection with no introgression and Neolithic migration models.

| | **B** | **M** | **S** |
|---|---|---|---|
| **B** | 100.00% | 0.00% | 0.00% |
| **M** | 0.00% | 100.00% | 0.00% |
| **S** | 0.00% | 0.00% | 100.00% |
| **Posterior model probabilities** | 0.00% | 100.00% | 0.00% |

Confusion matrix for misclassification is reported here using SMC for random samples from the models. Posterior model probabilities are final posterior after using the real data.

#### Supplementary Table 8: Cross validation and Model Selection with all the models together.

| | **BND** | **BNDF** | **BNDX** | **BNDXF** | **BNI** | **MND** | **MNDF** | **MNDX** | **MNDXF** | **MNI** | **SND** | **SNDF** | **SNDX** | **SNDXF** | **SNI** |
|---|---|---|---|---|---|---|---|---|---|---|---|---|---|---|---|
| **BND** | 71.0% | 15.4% | 9.3% | 4.2% | 0.0% | 0.0% | 0.0% | 0.0% | 0.0% | 0.0% | 0.0% | 0.0% | 0.0% | 0.0% | 0.0% |
| **BNDF** | 19.1% | 65.3% | 5.7% | 9.0% | 0.0% | 0.0% | 0.7% | 0.0% | 0.0% | 0.0% | 0.0% | 0.2% | 0.0% | 0.0% | 0.0% |
| **BNDX** | 7.0% | 2.6% | 65.3% | 25.0% | 0.0% | 0.0% | 0.0% | 0.1% | 0.0% | 0.0% | 0.0% | 0.0% | 0.0% | 0.0% | 0.0% |
| **BNDXF** | 3.3% | 8.1% | 20.1% | 68.0% | 0.0% | 0.0% | 0.0% | 0.0% | 0.4% | 0.0% | 0.0% | 0.0% | 0.0% | 0.1% | 0.0% |
| **BNI** | 0.0% | 0.0% | 0.0% | 0.0% | 100.0% | 0.0% | 0.0% | 0.0% | 0.0% | 0.0% | 0.0% | 0.0% | 0.0% | 0.0% | 0.0% |
| **MND** | 0.0% | 0.0% | 0.0% | 0.0% | 0.0% | 99.9% | 0.0% | 0.0% | 0.0% | 0.0% | 0.0% | 0.0% | 0.0% | 0.0% | 0.0% |
| **MNDF** | 0.0% | 0.9% | 0.0% | 0.0% | 0.0% | 0.0% | 96.7% | 0.0% | 0.5% | 0.0% | 0.0% | 1.8% | 0.0% | 0.0% | 0.0% |
| **MNDX** | 0.0% | 0.0% | 0.0% | 0.0% | 0.0% | 0.0% | 0.0% | 99.5% | 0.5% | 0.0% | 0.0% | 0.0% | 0.0% | 0.0% | 0.0% |
| **MNDXF** | 0.0% | 0.1% | 0.0% | 0.0% | 0.0% | 0.0% | 0.2% | 0.7% | 99.1% | 0.0% | 0.0% | 0.0% | 0.0% | 0.0% | 0.0% |
| **MNI** | 0.0% | 0.0% | 0.0% | 0.0% | 0.0% | 0.0% | 0.0% | 0.0% | 0.0% | 100.0% | 0.0% | 0.0% | 0.0% | 0.0% | 0.0% |
| **SND** | 0.0% | 0.0% | 0.0% | 0.0% | 0.0% | 0.0% | 0.0% | 0.0% | 0.0% | 0.0% | 100.0% | 0.0% | 0.0% | 0.0% | 0.0% |
| **SNDF** | 0.0% | 0.0% | 0.0% | 0.0% | 0.0% | 0.0% | 1.3% | 0.0% | 0.0% | 0.0% | 0.0% | 98.7% | 0.0% | 0.0% | 0.0% |
| **SNDX** | 0.0% | 0.0% | 0.0% | 0.0% | 0.0% | 0.0% | 0.0% | 0.0% | 0.0% | 0.0% | 0.0% | 0.0% | 100.0% | 0.0% | 0.0% |
| **SNDXF** | 0.0% | 0.0% | 0.0% | 0.0% | 0.0% | 0.0% | 0.0% | 0.0% | 0.2% | 0.0% | 0.0% | 0.0% | 0.0% | 99.8% | 0.0% |
| **SNI** | 0.0% | 0.0% | 0.0% | 0.0% | 0.0% | 0.0% | 0.0% | 0.0% | 0.0% | 0.0% | 0.0% | 0.0% | 0.0% | 0.0% | 100.0% |
| **Posterior model probabilities** | 0.0% | 0.0% | 24.3% | 75.7% | 0.0% | 0.0% | 0.0% | 0.0% | 0.0% | 0.0% | 0.0% | 0.0% | 0.0% | 0.0% | 0.0% |

Confusion matrix for misclassification is reported here using SMC for random samples from the models. Posterior model probabilities are final posterior after using the real data. ([B, M, S]x [No Introgression; Neanderthal and Denisova Introgression (ND); Neanderthal, Denisova and Africa archaic introgression (NDX); Neanderthal, Denisova introgression and Farming Migration (NDF); Neanderthal, Denisova, African archaic introgression and Farming Migration (NDXF)]

Supplementary Table 9: Posterior range of model B by ABC-DLS without using SMC.

| Parameters | Mean | CI |
|---|---:|---|
| **N_A** | 15,628 | 11,963 - 23,977 |
| **N_AF** | 48,024 | 16,870 - 123,355 |
| **N_EU** | 60,378 | 22,402 - 122,695 |
| **N_AS** | 84,417 | 30,100 - 142,309 |
| **N_F** | 17,439 | 5,847 - 29,281 |
| **N_EU0** | 3,109 | 1,997 - 4,798 |
| **N_AS0** | 744 | 519 - 1,042 |
| **N_BC** | 17,106 | 3,033 - 28,253 |
| **N_B** | 2,963 | 2,537 - 3,318 |
| **N_AF** | 15,313.00 | 4,469 - 27,566 |
| **T_FM (ky)** | 3.51 | 2.12 - 4.94 |
| **T_FS (ky)** | 4.64 | 0 - 9.31 |
| **T_DM (ky)** | 18.93 | 10.16 - 28.23 |
| **T_EU_AS (ky)** | 20.29 | 10.28 - 29.73 |
| **T_NM (ky)** | 14.52 | 9.47 - 19.63 |
| **T_XM (ky)** | 31.49 | 8 - 48.4 |
| **T_Mix (ky)** | 15.18 | 6.25 - 26.89 |
| **T_Sep (ky)** | 15.06 | 5.32 - 28.16 |
| **T_B (ky)** | 17.73 | 4.49 - 44.65 |
| **T_AF (ky)** | 284.27 | 0 - 680.92 |
| **T_N_D (ky)** | 412.26 | 338.49 - 448.3 |
| **T_H_A (ky)** | 209.6 | 131.82 - 246.93 |
| **T_H_X (ky)** | 566.81 | 449.16 - 684.47 |
| **Mix (%)** | 47.54 | 23.4 - 73.04 |
| **NMix (%)** | 2.88 | 2.65 - 3 |
| **DMix (%)** | 0.7 | 0.3 - 1.24 |
| **XMix (%)** | 2.3 | 0.05 - 9.09 |
| **FMix (%)** | 1.91 | 0.41 - 3.78 |

CI is the confidence interval of 2.5%-97.5% for respective parameters. Ky means kilo or thousand years.

#### Supplementary Table 10: Posterior range of model B using ABC-RF.

| Parameters | Expectation | CI |
|---|---|---|
| **N_A** | 15,706 | 11,098 - 22,943 |
| **N_AF** | 36,422 | 13,044 - 123,459 |
| **N_EU** | 68,755 | 17,388 - 141,245 |
| **N_AS** | 86,992 | 18,063 - 145,812 |
| **N_F** | 17,551 | 5,558 - 29,535 |
| **N_EU0** | 3,123 | 1,457 - 4,874 |
| **N_AS0** | 1,089 | 546 - 2,595 |
| **N_BC** | 15,063 | 1,642 - 29,243 |
| **N_B** | 3,904 | 2,417 - 4,952 |
| **N_AF0** | 19,098 | 7,461 - 29,145 |
| **T_FM (ky)** | 3.5 | 2.1 - 5 |
| **T_FS (ky)** | 5 | 0.4 - 9.7 |
| **T_DM (ky)** | 19.1 | 10.2 - 34.3 |
| **T_EU_AS (ky)** | 17.7 | 10.3 - 29 |
| **T_NM(ky)** | 17.8 | 5.6 - 42.2 |
| **T_XM (ky)** | 28.8 | 6.1 - 49.1 |
| **T_Mix (ky)** | 24.1 | 6.2 - 47.8 |
| **T_Sep (ky)** | 26.8 | 6.4 - 48 |
| **T_B (ky)** | 28.9 | 5.7 - 77.1 |
| **T_AF (ky)** | 340.7 | 14.1 - 688.4 |
| **T_N_D (ky)** | 393.1 | 333.2 - 447.2 |
| **T_H_A (ky)** | 192.3 | 124.2 - 247.9 |
| **T_H_X (ky)** | 575.2 | 458 - 694.2 |
| **Mix (%)** | 54.42 | 8.16 - 93.2 |
| **NMix (%)** | 2.28 | 1.11 - 2.98 |
| **DMix (%)** | 0.86 | 0.08 - 1.89 |
| **XMix (%)** | 5.07 | 0.41 - 9.76 |
| **FMix (%)** | 4.19 | 0.24 - 9.51 |

CI is the confidence interval of 2.5%-97.5% of respective parameters. Ky means kilo or thousand years.

Supplementary Table 11: Posterior range for parameters of model B with slower mutation rate.

| Parameters | Mean | CI | Events (kya) |
|---|---|---|---|
| **N_A** | 15,792 | 15,720 - 15,931 | |
| **N_AF** | 22,719 | 22,376 - 23,204 | |
| **N_EU** | 77,929 | 76,751 - 80,340 | |
| **N_AS** | 124,981 | 122,317 - 129,200 | |
| **N_F** | 34,288 | 33,804 - 34,834 | |
| **N_EU0** | 2,618 | 2,608 - 2,636 | |
| **N_AS0** | 874 | 868 - 886 | |
| **N_BC** | 17,843 | 16,903 - 21,127 | |
| **N_B** | 2,385 | 2,366 - 2,401 | |
| **N_AF0** | 28,376 | 28,156 - 28,819 | |
| **T_FM** | 3.3 | 1.9 - 4.4 | 3.3 (1.9 - 4.4) |
| **T_FS** | 5.5 | 1.2 - 9.6 | 8.7 (4.4 - 13.1) |
| **T_DM (ky)** | 14.8 | 14.6 - 15.1 | 14.8 (14.6 - 15.1) |
| **T_EU_AS (ky)** | 23 | 22.8 - 23.3 | 37.8 (37.4 - 38.2) |
| **T_NM (ky)** | 5.9 | 5.5 - 6.1 | 43.7 (43.2 - 44.2) |
| **T_XM (ky)** | 15.3 | 13.8 - 17.9 | 53.1 (51 - 55.2) |
| **T_Mix (ky)** | 21.6 | 21.2 - 22 | 59.4 (58.8 - 59.9) |
| **T_Sep (ky)** | 7.6 | 7.3 - 7.9 | 67 (66.3 - 67.6) |
| **T_B (ky)** | 13.7 | 13.5 - 14 | 80.6 (80 - 81.3) |
| **T_AF (ky)** | 251 | 250.4 - 252.3 | 331.6 (330.4 - 332.8) |
| **T_N_D (ky)** | 439.2 | 434.5 - 448 | 439.2 (434.5 - 448) |
| **T_H_A (ky)** | 239.2 | 234 - 246.2 | 678.5 (669.4 - 687.6) |
| **T_H_X (ky)** | 684.8 | 681.1 - 692.7 | 684.8 (681.1 - 692.7) |
| **Mix (%)** | 90.13 | 89.09 - 90.63 | |
| **NMix (%)** | 2.91 | 2.83 -2.99 | |
| **DMix (%)** | 0.71 | 0.67 - 0.78 | |
| **XMix (%)** | 6.57 | 6.5 - 6.73 | |
| **FMix (%)** | 1.9 | 1.87 - 2 | |

CI is the confidence interval of 2.5%-97.5% of respective parameters. Ky means kilo years and kya means kilo or thousand years ago from now.

#### Supplementary Table 12: Relations of events with the time intervals that were used in the simulations

| **Events** | **Model S** | **Model B** | **Model M** |
|---|---|---|---|
| **Farming Admixture** | T_FM | T_FM | T_FM |
| **Farming Separation (FS)** | T_FM + T_FS | T_FM + T_FS | T_FM + T_FS |
| **Denisova Introgression** | T_DM | T_DM | T_DM |
| **Split Europe Asia (E_A)** | max (T_DM, FS) + T_EU_AS | max (T_DM, FS) + T_EU_AS | max (T_DM, FS) + T_EU_AS |
| **Neanderthal Introgression (NI)** | E_A + T_NM | E_A + T_NM | E_A +T_NM |
| **African Introgression (AI)** | E_A + T_XM | E_A + T_XM | E_A + T_XM |
| **Admixture (Mix)** | NA | E_A + T_Mix | E_A + T_Mix |
| **Separation (Sep)** | NA | Mix + T_Sep | Mix + T_Sep |
| **Split Africa OOA (OOA)** | max (NI, AI) + T_B | max (NI, AI, Sep) + T_B | max (NI, AI, Sep) + T_B |
| **Ancestral Size change** | OOA + T_AF | OOA + T_AF | OOA + T_AF |
| **Split Neanderthal Denisova** | T_N_D | T_N_D | T_N_D |
| **Split Human Neanderthal** | T_N_D + T_H_A | T_N_D + T_H_A | T_N_D + T_H_A |
| **Split human African archaic** | T_H_X | T_H_X | T_H_X |
